# Comprehensive characterization of genomic, transcriptomic and epigenomic artifacts introduced in formalin-fixed, paraffin-embedded tissues

**DOI:** 10.64898/2026.08.31.748145

**Authors:** Erik J Zmuda, Kyle R Covington, Cyriac Kandoth, Charlotte K Y Ng, Daniel J Weisenberger, Reanne Bowlby, Andy Chu, Richard Corbett, Denise Brooks, Andrew D Cherniack, Bradley Murray, Shiyun Ling, Wei Zhao, Eve Shinbrot, Raymond S Lim, Moiz S Bootwalla, Toshinori Hinoue, Jianhong Hu, Nipun Kakkar, Michael Lawrence, Michael McLellan, Christopher Miller, Donna Morton, Andrew J Mungall, Donna Muzny, Carrie Sougnez, Petar Stojanov, Liu Xi, Nicholas Schultz, Rehan Akbani, Christina Curtis, Bob Fulton, Richard A Gibbs, Gad Getz, Paul Spellman, Marco Marra, Marc Ladanyi, Roy Tarnuzzer, Heidi Sofia, the Cancer Genome Atlas Research Network, Carolyn Hutter, David A Wheeler, Julie Gastier-Foster, Peter W Laird, HarshaVardhan Doddapaneni, Jorge S Reis-Filho, Jean Claude Zenklusen, Katherine A Hoadley

## Abstract

Genomic, transcriptomic and epigenomic characterization has accelerated the discovery of clinically-relevant alterations in cancer, predominantly using fresh frozen (FF) specimens. However, clinical molecular pathology laboratories prefer formalin-fixed paraffin-embedded (FFPE) methods, known to introduce artifacts at the nucleic acid level, over fresh frozen methods. Extending the multi-platform analysis to FFPE specimens for comprehensive clinical molecular diagnosis requires a thorough understanding of the consequence of formalin-fixation. We present a detailed multi-platform characterization of FFPE preservation using paired FF specimens as the ’gold standard’. DNA and RNA were obtained from 38 patients across 6 cancer types using a FFPE optimized co-isolation. The impact of FFPE on exome sequencing was dependent on filtering, where a minimum coverage or supporting read filter can mitigate FFPE-specific false positives. Copy number alterations, MSI assessment, mutational signatures, and DNA methylation were comparable between FFPE and FF. FFPE biases in RNA expression can be overcome when using biology-relevant genes and we describe a novel consequence of FFPE on miRNA species diversity. Collectively, this data provides a broad view of FFPE artifact and offers best practices for overcome these biases.

## INTRODUCTION

Massively parallel sequencing (MPS) and array based methods to characterize the landscape of genomic, transcriptomic and epigenomic alterations in cancer have resulted in major advancements in our understanding of tumor biology (2015a; 2015b; Hoadley et al.; The Cancer Genome Atlas, 2015). Diagnostic tests are now utilizing these platforms to detect clinically relevant alterations (Duncavage et al., 2012; Ellis et al., 2012; Ley et al., 2010; Walter et al., 2012). Clinical molecular pathology laboratories prefer formalin-fixed paraffin-embedded (FFPE) methods over fresh frozen (FF) preservation since FFPE produces higher quality histological images and is easier to store. Unfortunately, formalin fixation is well known to introduce artifacts at the nucleic acid level. Formaldehyde reacts at the molecular level by introducing covalent methylene bridges between DNA-DNA, DNA-RNA, and DNA-Protein. It also induces oxidative and deamination reactions in both DNA and RNA that ultimately result in the formation of cyclic base derivatives (Auerbach et al., 1977; Farragher et al., 2008b; Feldman, 1973; Karlsen et al., 1994; Srinivasan et al., 2002). These modifications invariably complicate molecular diagnostic testing and data interpretation because they lead to nucleic acid fragmentation, base substitutions and inhibit enzymatic reactions. FFPE-specific workflows, including methods for nucleic acid extraction, wet-lab protocols, and bioinformatics/analytical procedures are required to counteract the consequences of formalin fixation and to separate analytical artifacts from true biological events.

Numerous studies have characterized the spectrum of molecular consequences of FFPE and/or evaluated the feasibility of clinical molecular diagnostics using FFPE-derived analytes including DNA (Do and Dobrovic, 2012; Do et al., 2013; Hedegaard et al., 2014; Kerick et al., 2011; Schweiger et al., 2009; Spencer et al., 2013; Van Allen et al., 2014; Wagle et al., 2012; Wong et al., 2014; Yost et al., 2012), RNA (April et al., 2009; Farragher et al., 2008a; Hedegaard et al., 2014; Kibriya et al., 2010; Norton et al., 2013; Zhao et al., 2014), miRNA (Lehmann, 2010; Lim et al., 2015; Meng et al., 2013; Zhang et al., 2008), and DNA methylation (Conway et al., 2011; Moran et al., 2014). Overall, these studies indicate that FFPE can be utilized for specific clinical tests; however, results are not always equivalent to those obtained from FF. Differences, such as low-allele fraction single-base substitutions in DNA or non-reproducible detection of RNA fusions, introduce the potential for inaccurate diagnostic reporting. While these studies have deepened our understanding of FFPE artifacts, they are individually associated with unique limitations, which include the absence of paired FF tissue, narrowly targeted characterization (gene panels versus whole exome or genome sequencing), single platform analysis, or insufficient descriptive granularity regarding the artifact introduced by FFPE preservation thus hindering the development of effective bioinformatic filters.

To increase the diagnostic utility of FFPE-preserved specimens to a broader community of users, an in-depth comparison of paired FF and FFPE tissues across multiple platforms is needed. In this study we present a detailed molecular signature of FFPE preservation, identified by using paired FF specimens as the ‘gold standard’ for analyzing results from whole exome and whole genome sequencing, SNP6.0 array copy number profiling, total RNA and miRNA sequencing, and Illumina Infinium HM450 DNA methylation profiling. Results are presented for 38 patients, representing 6 tumor types.

## RESULTS

### Sample processing, analyte extraction and quality control

We employed a RNA and DNA co-isolation extraction protocol to maximize analyte integrity. Paired FF and FFPE-preserved control thymus tissues were used for protocol optimization with the initial quality of each tissue determined from the FF portion. No single commercial co-isolation protocol following manufacturer’s recommendations was able to successfully produce optimal quality of both DNA and RNA; some methods preserved DNA integrity at the expense of the RNA and vice versa (Supplementary Figure 1a). Merging components from different kits with optimization of reagent volumes led to a protocol that yielded maximal integrity of co-isolated DNA and RNA (Supplementary Figure 1a,b).

Using our optimized extraction protocol, we co-isolated DNA and RNA from FFPE tumor biospecimens. All FFPE-derived analytes were paired with analytes extracted from an adjacent FF portion of the same tumor biospecimen, as well as blood germline DNA. This study included 38 patients from 6 tumor types, including bladder urothelial carcinoma (BLCA; n=3), endometrial carcinoma (UCEC; n=4), kidney renal clear cell carcinoma (KIRC; n=4), invasive breast carcinoma (BRCA; n=5), colon adenocarcinoma (COAD; n=10) and lung adenocarcinoma (LUAD; n=12). To quantify the extent of tumor heterogeneity associated with the spatial separation of paired FF and FFPE specimens, additional adjacent portions of FF tumors from seven LUAD patients were extracted and included in this study. A description of specimens used, the amount of material required for extraction and the quality control metrics from the extracted analytes are provided in Supplementary Table 1 and Supplementary Figure 1c. Analytes were distributed for whole exome sequencing, whole genome sequencing, SNP6.0 copy number profiling, total RNA and miRNA sequencing, miRNA sequencing and DNA methylation analysis (Supplementary Figure 1d)

### Whole Exome Sequencing (WES)

WES was performed on paired tumor (FF and FFPE) and germline normal genomic DNA from 38 tumors at one of three large scale sequencing centers identified as Pipeline 1, Pipeline 2 and Pipeline 3. Somatic single nucleotide variants (SNVs) and insertions/deletions (INDELs) detected in tumors were quantified using non-reference base calling (nREF) and Multi-center calling (MCC) as described in the methods.

We observed a surprisingly low overall rate of concordance between FF and FFPE WES data (Supplementary Table 2). In the nREF call set, the concordance was 33.47% for SNVs and 66.63% for INDELs while in MCC the union was 22.48% for SNVs and 36.59% for INDELs and only slightly improved when restricted to variants detected by two or more pipelines (38.4% SNVs; 44.86% INDELs). Using variant allele fraction (VAF) to create bins, we found that the overwhelming majority of FFPE-unique events (nREF: 96.74% SNVs, 88.14% INDELs; MCC: Union: 93.3% SNVs, 70.89% INDELs; Intersection: 92.08% SNVs, 62.5% INDELs) resided in the <20% VAF (Figure 1a and 1b). Individual variant calling pipelines had a large impact on FFPE-unique variants; Pipeline 3 gave rise to minimal FFPE-unique events while Pipelines 1 and 2 retained a large number of FFPE-unique events in the <20% VAF (Figure 1b).

**Figure 1.**
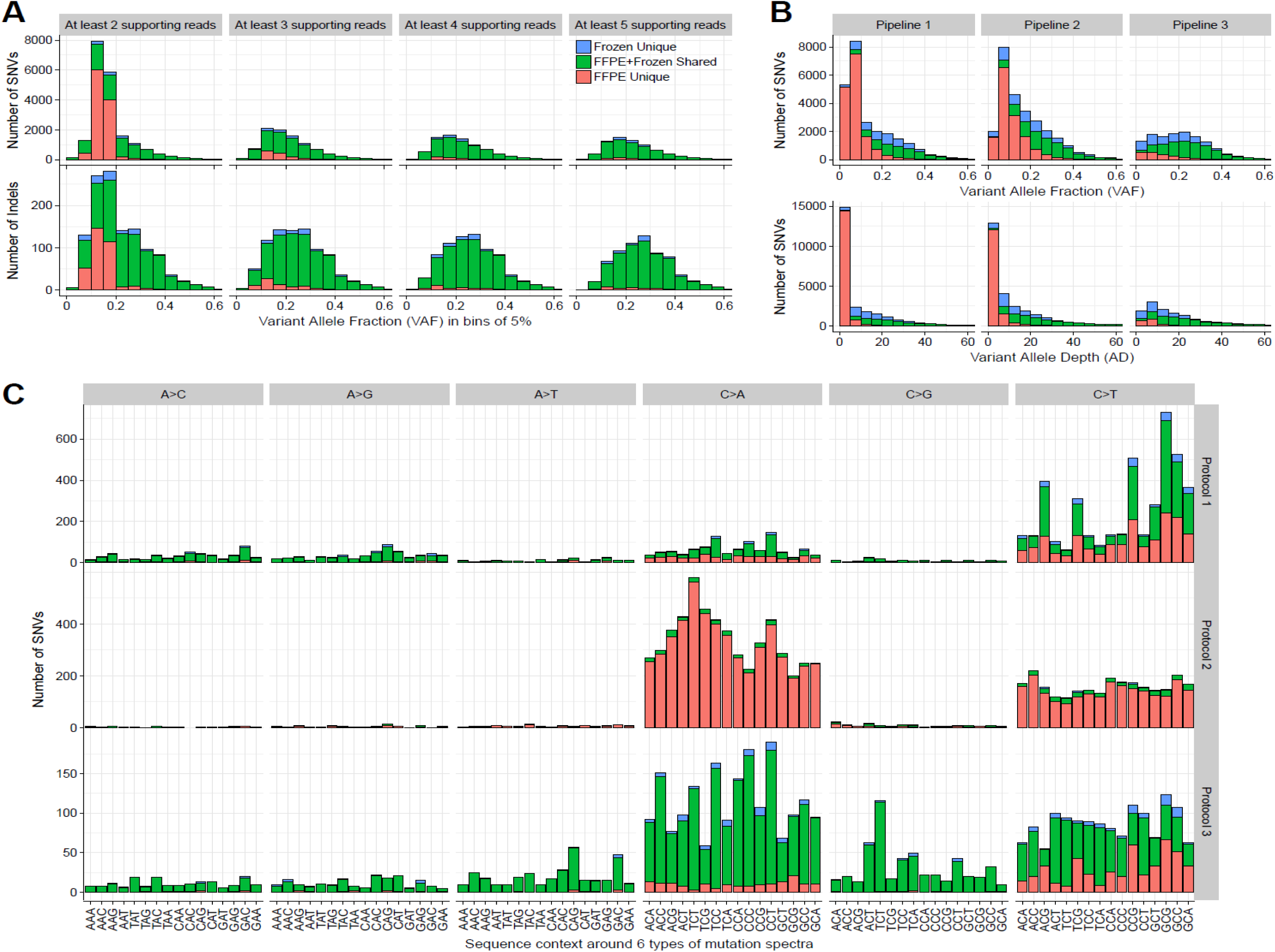
The mutational signature of FFPE artifact as viewed through either a minimally filtered variant calling pipeline or multiple fully optimized pipelines. (a) Frequency and allele fraction of all SBS and INDELs found unique to FFPE (Salmon), shared between FF and FFPE (Green), or unique to frozen samples (Blue) from non-reference base calling stratified by minimum supporting reads. (b) Allele fraction and allele depth as shown in panel (a) for variants called by individual optimized pipelines. (c) Sequence context of nREF distribution for all combinations of SBS generated by protocols in place at individual sequencing facilities.

By increasing the minimum number of supporting reads within nREF, we could decrease the frequency of FFPE-unique events with ≥5 reads eliminating 97.5% of all FFPE-unique events (Figure 1a, Supplementary Table 2). However, this filter also excluded 37.4% of variants detected in both FF and FFPE (presumed true positives), highlighting the need to balance specificity with sensitivity. Similarly, this filter would also be effective in Pipeline 1 and 2 as a majority of FFPE-unique events are restricted to the ≤5 supporting read bin (Figure 1b). Pipeline 3 would be relatively unaffected by this filter since it had few FFPE-unique called events (Figure 1b).

In the nREF call set, variants unique to FFPE-preserved specimens in the low VAF were predominately comprised of single base substitutions (SBS) including C>T transitions (classic FFPE signature) and C>A transversions (Figure 1c). While C>A transversions were associated with a specific sequence context (N[C>A][A/T]), we did not detect a strong association between sequence context and C>T transitions (Supplementary Figure 2) except for KIRC and COAD, where C>T transitions were associated with N[C>T]G. Most FFPE-unique SBS detected by nREF were filtered out by requiring a minimum of five supporting reads or a ≥ 20x sequencing depth coverage (Supplementary Figure 2). BRCA and UCEC contained a greater proportion of FFPE-unique events across both low (<20%) and high (>20%) VAF than other tissues (Supplementary Figure 2) and neither the minimum supporting read or total read depth filter was effective at mitigating the effect of FFPE. These BRCA and UCEC samples were sequenced at only one center; therefore, it is not possible to determine if this is due to a specific FFPE artifact, the tumor type, or the sequencing center.

### Detecting Biologically Significant Somatic Alterations from WES of FFPE Tissue

We focused on a subset of significantly mutated genes (SMG) previously identified in TCGA and evaluated the concordance between FFPE and FF in MCC (Figure 2a). Despite the overall low concordance observed between FF and FFPE, the majority of variants in SMGs were found in both (100/131, 76%). When detected, FFPE-unique events were predominately C>A transversions in BRCA and UCEC samples and most (44/52; 85%) had < 5 supporting reads and VAFs below minimal thresholds (Figure 2b). FF-unique mutations were also detected in 4 out of 6 tumor types, across a range of SBSs and VAFs. To our surprise, a manual review of the corresponding paired FFPE BAM files for these FF-unique events did confirm the presence of supporting reads in the FFPE sample, indicating that the variant was present but failed to be called by any of the mutation calling pipelines. No explanation could be found to account for the absence of these variants from multiple pipelines’ variant call files. Thus, the signature of FFPE artifacts is not limited to the introduction of novel base errors, but can also involve a loss in sensitivity for detecting potentially true variants.

**Figure 2.**
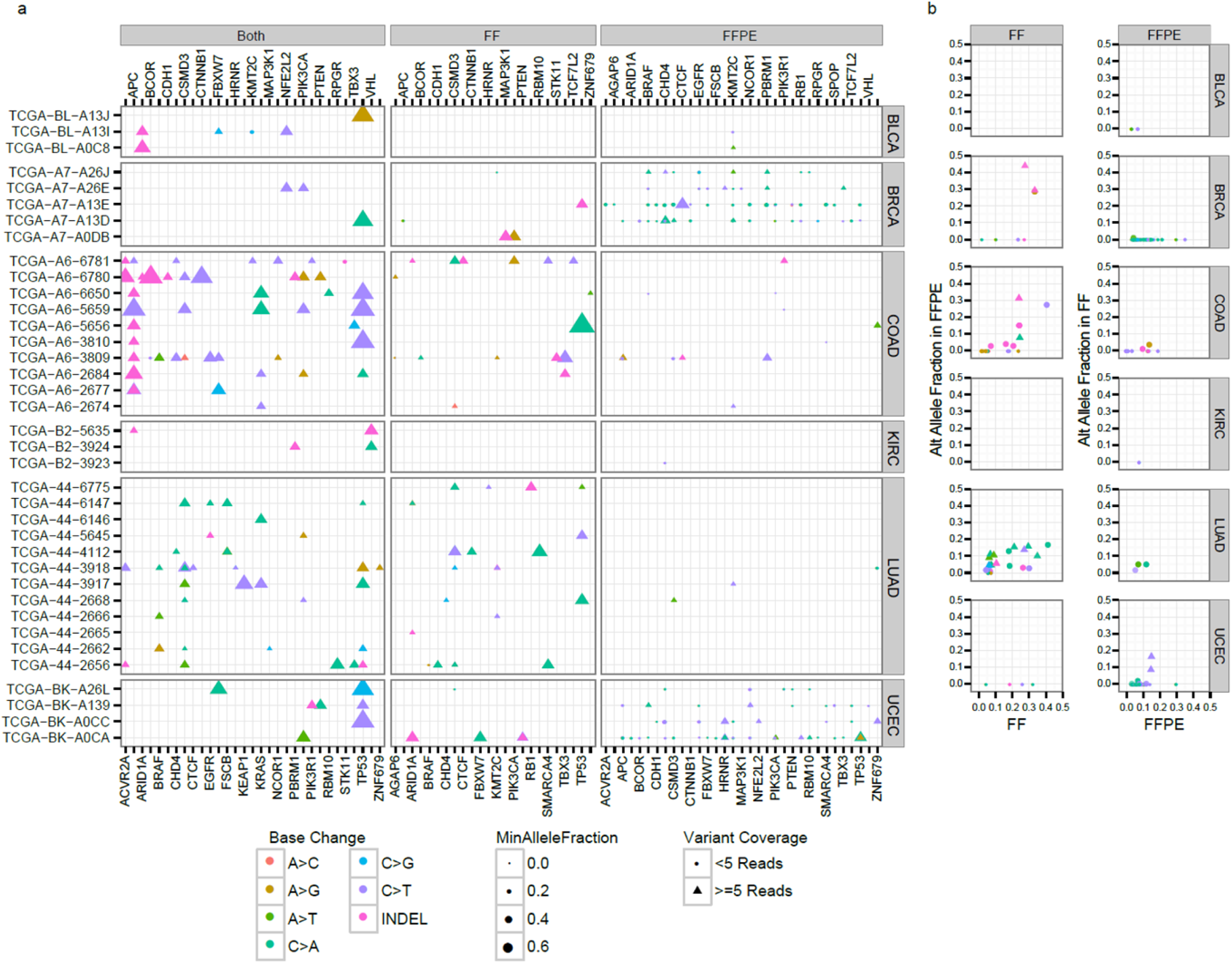
Detection of biologically significant events in FFPE tumors. (a) Dot-plot of all variants called by large scale sequencing centers in cancer genes for all studies (major rows) and different detection modes (major columns, Both = detected in all preservation methods, FF = detected in FF only, FFPE = detected in FFPE only). Subjects and genes are represented as rows and columns respectively. Symbol shape represents the total number of reads supporting the variant (triangle = 5 or more, circle = less than 5), symbol size represents the allele fraction for the call and symbol color indicates base change. (b) Dot-plot of unique variants in the discovery preservation method (major columns) and alternate preservation method for all studies (major rows). Discovery allele fraction and allele fraction in the alternate preservation method after lookup are shown in the x and y-axis respectively.

Microsatellite instability (MSI) is found in approximately 15% of colorectal cancers and 30% of endometrial cancers (Boland and Goel, 2010; Kandoth et al., 2013). We evaluated the detection of MSI in COAD and UCEC samples from this cohort. We calculated the MSI homopolymer scores from WES data as previously described (Shinbrot et al., 2014) and compared to the MSI status determined previously from patient matched FF samples by the gold standard mono/di-nucleotide repeat PCR assay (Parikh et al., 2014; Walter et al., 2012). The homopolymer scores between FF and FFPE samples were concordant (Supplementary Figure 3). The three COAD patients (A6-3809, A6-6780, and A6-6781) classified as MSI-High using the PCR assay had the highest homopolymer scores. These results show that FF and FFPE WES data can yield comparable MSI results.

Mutational signatures can help understand tumorigenesis and the mutational process in cancer. However, FFPE-induced artifact may compromise the accuracy of signature analysis. To evaluate the impact of FFPE, we first used a non-smooth non-negative matrix factorization (nsNMF) method (BioRxiv: https://doi.org/10.1101/036541) to analyze the mutational signatures in the FF specimens (Supplementary Figure 4a). We found that Signatures 1 (AC>AN;AT>AN), 4 (*APOBEC*, BLCA), 5 (smoking, LUAD), 6 (CpG, COAD, UCEC, BRCA) and 12 (Not Defined) were enriched in our FF cohort and consistent with the tissue types assayed (Supplementary Figure 4a). The Signatures were highly similar between FF and FFPE samples, except for Signature 1, evidenced by a highly significant correlation between signature coefficients in FF and FFPE (Supplementary Figure 4b). These results confirm that FFPE preservation does not significantly alter the overall mutational signatures detected through nsNMF.

We used the same approach to understand if FF- and FFPE-unique variants were related to formalin fixation or tumor spatial heterogeneity. Using the additional FF portion for seven LUAD tumors, we quantified sequence-context specific low VAF (<20%) SBS in paired FF portions and FF-FFPE pairs separately (Supplementary Figure 5a). In both, there was an enrichment of C>T transitions; however the sequence context did not strongly overlap. To help distinguish the signature of spatial heterogeneity within multiple portions of the same tumor from artifact introduced by FFPE, we further separated variants into those that were shared between paired FF portions (heterogeneity-shared), those that were unique between the paired FF portions (heterogeneity-unique) and those that were unique between paired FF-FFPE portions (FFPE-unique). Similar to the full analysis, Signatures 1, 5, 6 and 12 were the most penetrant. The heterogeneity-shared variants contributed to Signature 5 (smoking) and were observed within both high and low VAF (Supplementary Figure 5b). FFPE-unique events were predominantly low VAF (<20%) and enriched in Signature 6 (CpG), consistent with C>T transitions observed in FFPE (Supplementary Figure 5b and 5c). The heterogeneity-unique events displayed similar features with predominantly low VAF and a Signature 6 profile, but with substantially less intensity than that observed from FFPE-unique events (Supplementary Figure 5c and 5d). These results suggest that while FFPE-unique events represent a combination of variables introduced by formalin fixation and tumor spatial heterogeneity, a significant majority of the FFPE artifact we report from WES stems from formalin fixation.

### Somatic Copy Number Alteration (SCNA) analysis

We used three approaches to compare genome-wide patterns of SCNAs in 27 FF and FFPE tumor pairs: Affymetrix SNP6 array, WES and WGS (Figure 3a). For SNP6, SCNA patterns seen in FF tumors were often absent or greatly attenuated in FFPE. In contrast, the SNCA patterns generated from WES or WGS more closely resembled each other. To quantify the impact of FFPE, we derived gene level copy number estimates and determined the correlation between paired FF and FFPE tumors (Fig 3b). As expected for a platform that was not designed for FFPE, SNP6 correlations were lower than those observed in sequencing data (median SNP6 r^2^ 0.46; median WES r^2^ 0.75; median WGS r^2^ 0.85). We investigated overall noise across methods by quantifying the level of segmentation (Figure 3c). FFPE SNP6 data showed significantly more segmentation than paired FF samples (median 348 for FFPE and 106.5 for FF, p < 10^-6^ paired t-test), which was further observed at the probe level (Supplementary Figure 6a and b). WES and WGS segmentation values did not differ significantly between FF and FFPE tumors (WES median 223.5 for FF and 191.5 FFPE; WGS median 1057 for FF and 1736 for FFPE); however, the overall number of segments in both FF and FFPE WGS data was higher than that observed from either SNP6 or WES data. Despite the already high gene level copy number correlation, this suggests that additional preprocessing could further improve copy number analysis results from WGS. It is important to note that the copy number pipeline used in this study was designed to utilize WES data, and was not optimized for WGS. Given, that probe level noise data for WGS was less than that for SNP6 (Supplementary Figure 6a and b), it is plausible that algorithmic improvements could overcome segmentation issues in both FF and FFPE WGS SCNA data, making WGS more suitable for SCNA analysis.

**Figure 3.**
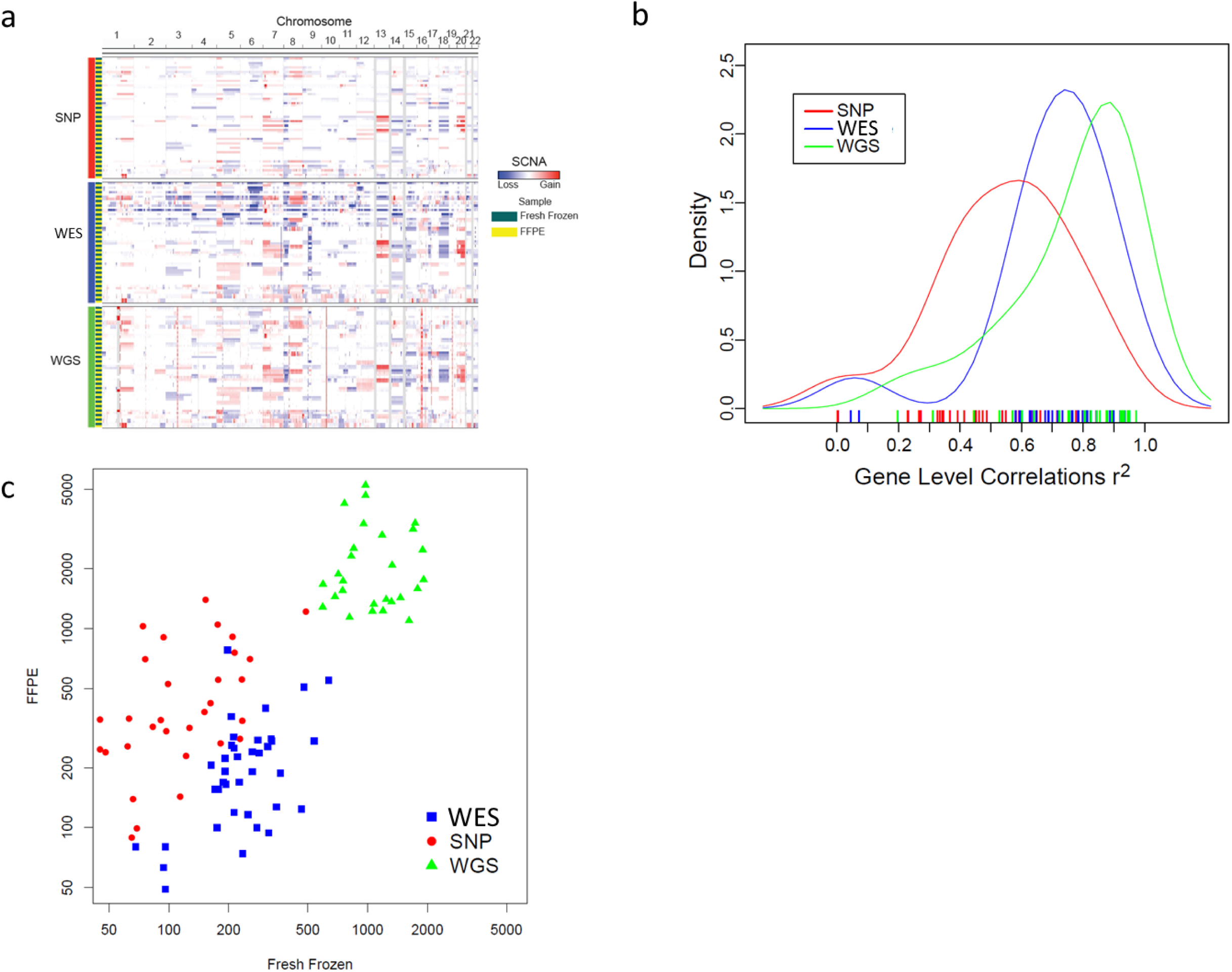
Analysis of somatic copy number alterations (SCNA) in FFPE tumors. (a) Landscape of SCNAs in FF and FFPE samples profiled by SNP6, WES and WGS analysis. In the heatmap, SCNAs in each tumor (rows) are plotted along genomic locations (columns). Red bars are regions of amplification and blue bars are regions of deletion. (b) Density plot of Pearson correlations of gene level copy number between FF and FFPE samples. (c) Comparison of copy number breaks (segments count) between FF and FFPE samples.

### RNA sequencing

Total RNAs from 38 FF and FFPE tumor pairs were sequenced using the Illumina Truseq Ribo Gold protocol to evaluate effectiveness of FFPE-derived RNA sequencing (Hedegaard et al., 2014; Zhao et al., 2014). While there were no differences in the total number of reads between FF and FFPE, the location of where reads mapped differed (Supplementary Figure 7a). Most notable was a decrease in reads mapping to coding and UTR regions and a corresponding increase in intronic reads in FFPE samples. We observed less consistent read distribution across genes in FFPE compared to FF, likely due to the strong 3’ bias in FFPE (Supplementary Figure 7b-d). The median insert size of FFPE libraries, predominately driven by LUAD and COAD samples, was significantly lower than that observed in FF (p=0.0004), a finding indicative of degraded RNA from FFPE (Supplementary Figure 7e).

Pairwise comparisons of transcript levels between FF and FFPE samples revealed high transcriptome-wide correlation (average r^2^>0.85) (Figure 4a and Supplementary Figure 8a). In a principal component analysis (PCA) of expressed genes (n=16,868), the first principle component was dominated by preservation status, while the second was tumor type (Supplementary Figure 8 b,c). Unsupervised clustering of 3,000 most variably expressed genes resulted in samples clustered by tissue type (Figure 4b). While UCEC and BLCA tumors clustered by patient, the remaining tissues clustered according to preservation method rather than patient (Figure 4b, 13/38 paired). We identified 1,802 genes associated with preservation method across all tumor types (q≤0, Figure 4c,d). COAD and LUAD tumors had the most differentially expressed genes between FF and FFPE [LUAD 1,390 (6.8% of genes); COAD 1,345 (6.6%); BRCA 194 (0.95%); KIRC 49 (0.24%); BLCA 3 (0.01%); and UCEC (0); q≤0] and were also the tumor types with the largest difference in quality control metrics. Clustering of the 3,000 most variable genes after removing the 1,802 preservation-associated genes improved clustering of samples by patient (Figure 4e, 36/38 paired). Unfortunately, excluding the same 1,802 genes from an independent cohort of paired FF and FFPE tumors did not reduce FFPE bias in unsupervised clustering (data not shown). While the external data set utilized RNA extracted and sequenced by the same methods, the age of the FFPE blocks were older and from a different tumor type (thyroid), adding to the complexity in developing signatures to help correct for FFPE biases. However, using biologically relevant gene lists derived from each tissue type ((2012c; 2014a; 2014b; Silver et al., 2013; Walter et al., 2012)), we were able to reduce the impact of preservation method for most tumor types. We performed paired T-tests on PC1 loadings between FF and FFPE samples and found the difference between pairs was reduced when using biology-specific genes (Supplementary Figure 9). Of note, if variable gene selection was also performed within FF or within FFPE, it also reduced FFPE bias (Supplementary Figure 9). These results indicate that RNA from FFPE samples can be used for RNA-sequencing and the results are highly consistent with RNA extracted from FF samples, particularly among genes with known biological importance.

**Figure 4.**
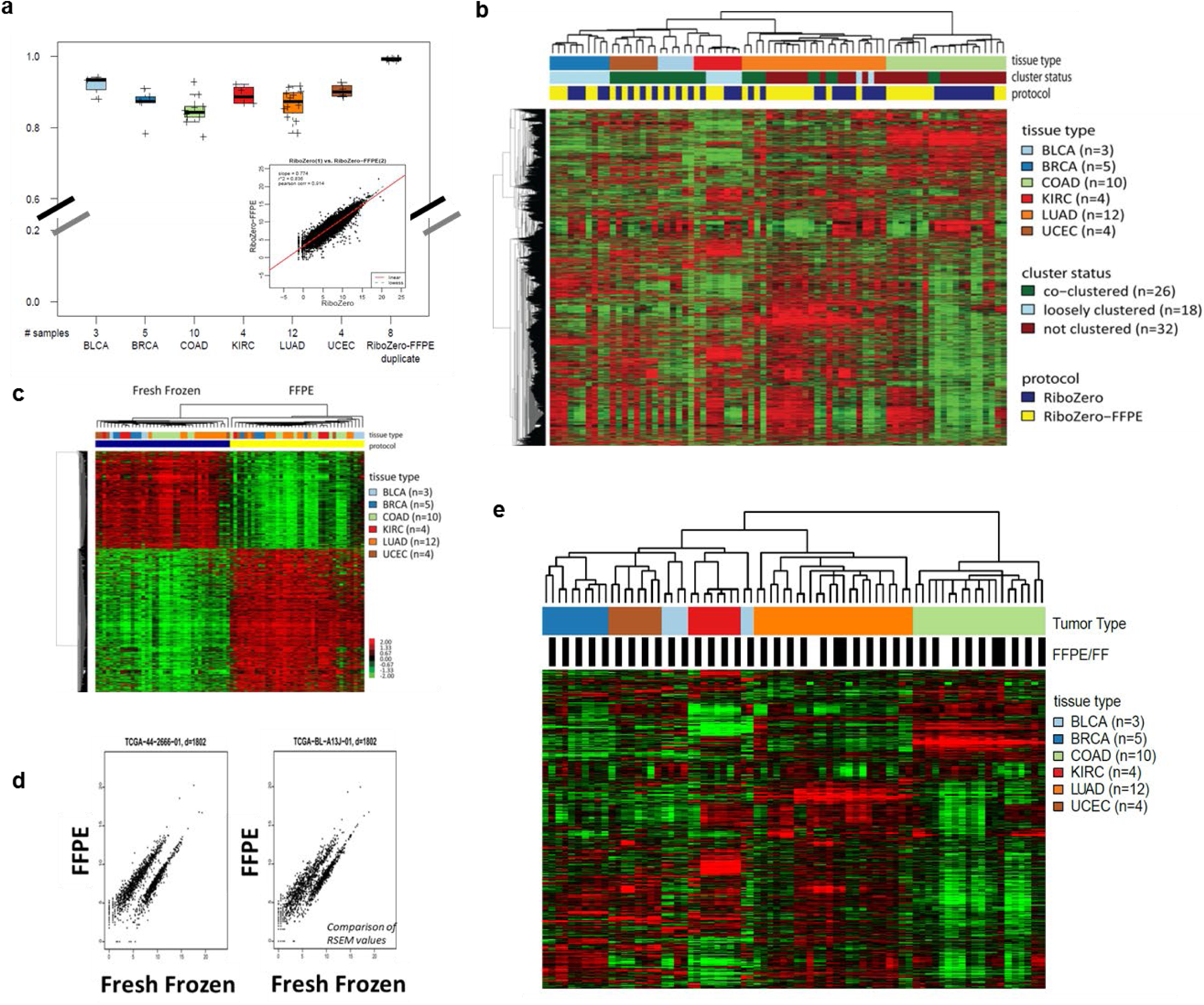
The signature of mRNA expression from FFPE tumors. (a) Pairwise Pearson correlation of transcript quantification between paired FF and FFPE tissues (inset shows representative patient). (b) Unsupervised hierarchical clustering of all samples using expression data from the 3,000 most variably expressed genes. (c) Clustering of significantly differentially expressed genes by preservation method. (d) Two representative examples of RSEM values for significantly differentially expressed transcripts. (e) Unsupervised clustering after removal of the top 1,802 differentially expressed genes.

### Detecting RNA transcript fusions from FFPE preserved tumors

We evaluated the concordance of fusion detection in FF and FFPE. A large number of novel fusions were detected in each of the samples; however, only one oncogenic recurrent fusion was observed (Supplementary Table 3). The recurrent in-frame fusion CTLC-ROS1 was detected in both LUAD TCGA- 44-2665 FF and FFPE samples and had been detected in a separate FF RNA aliquot previously sequenced in TCGA (2014b). No other recurrent oncogenic fusions were detected in our sample set or the previous TCGA marker papers ((2012c; 2014a; 2014b; Kandoth et al., 2013; Silver et al., 2013; Walter et al., 2012); cBioPortal). We are limited in our ability to comment on the feasibility of fusion detection in FFPE; however, having detected the single recurring oncogenic fusion in this cohort from both FF and FFPE supports the notion that valid fusions can be detected in FFPE preserved specimens.

### DNA Methylation analysis

We evaluated differences in DNA methylation between FF and FFPE tissues from 32 tumor pairs across 6 tumor types. We performed unsupervised hierarchical clustering using probes unmethylated in normal tissue samples corresponding to those included in this study (Figure 5a). The different preservation methods conserve sufficient biological information that unsupervised clustering organizes samples together not only by tissue type (32/32), but also by paired patient status (29/32 sample pairs). FF-FFPE sample pairs have extremely high correlation coefficients (r^2^≥0.95), validating that preservation method did not greatly differ in their effect on DNA methylation data (Figure 5b). We compared gene level DNA methylation profiles using *MLH1* as an example and found they were highly concordant (Figure 5c). However, we did identify discordant DNA methylation for one sample (TCGA-BK-A0CA). We evaluated CpG density distribution of the differences between the all of the FF and FFPE samples to look for sources of the discordance and identified a very slight skew towards higher DNA methylation values in the FFPE samples at lower CpG densities (Supplementary Figure 10a). This difference is very minor with a median differential DNA methylation beta value of -0.0168 (95% Confidence Interval: -0.0167, -0.0169) and can be attributed to the CpG density of the probes (Supplementary Figure 10b). Overall, FF and FFPE tissues generate highly comparable DNA methylation results.

We also evaluated epigenetic silencing and identified 216 epigenetically silenced genes in FF and 267 epigenetically silenced genes in FFPE. 115 genes were epigenetically silenced in both FF and FFPE (Supplementary Figure 11); demonstrating a trend towards decreased RNA expression in the presence of increased DNA methylation, while highlighting the variability that exists between FF and FFPE samples. We compared Z-scores for each platform and found that differences in gene expression levels between FF and FFPE had a greater influence than DNA methylation on the observed differences in silencing calls (Figure 5d). Overall, these results indicate that epigenetic silencing can be evaluated from FFPE derived analytes; however, their accuracy is limited by the cumulative effect of preservation method on both the RNA-seq and DNA methylation platforms.

### miRNA sequencing

We evaluated the differences in miRNA expression between FF and FFPE from 22 tumor pairs across 6 tumor types. miRNAs were successfully extracted and sequenced from the FFPE preserved specimens with some notable differences between preservation methods. The diversity of miRNA species identified in FFPE was slightly higher than FF (paired t-test p<0.03) despite a lower total number of aligned reads in FFPE (paired t-test p=0.02) (Supplementary Figure 12a,b).

Most pairs had a high Pearson correlation (rho > 0.92), with three exceptions (0.63-0.82) (Figure 6a). Similar to mRNA results, unsupervised hierarchical clustering using the top 25% most variable miRNAs resulted in a majority (30/44, 68%) of samples clustering by patient (including the three samples with low Pearson rho values)(Figure 6b). However, we identified 91 differentially expressed (DE) miRNAs between the two preservation methods with no clear quality metric that differentiated them from other reads (Figure 6c, Supplementary Table 4). Removing these DE miRNAs only slightly improved the number of samples that paired in clustering (32/44, Supplementary Figure 12c).

**Figure 5.**
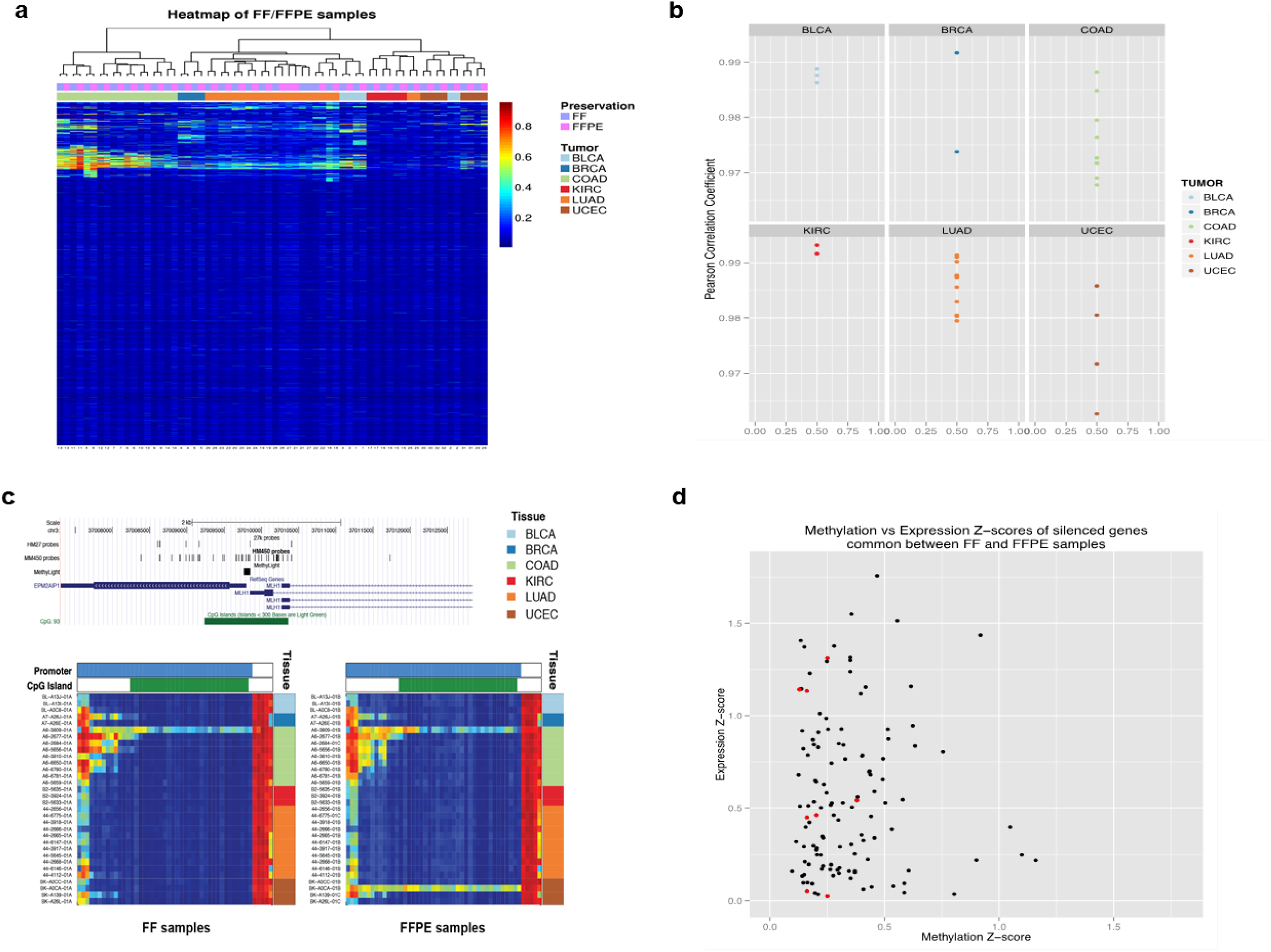
The signature of epigenetic DNA methylation and epigenetic silencing from FFPE tissues. (a) Heatmap visualization of DNA methylation from FF and FFPE samples after unsupervised, two-dimensional hierarchal clustering (32 FF-FFPE pairs (x-axis) at 12,849 CpG loci (y-axis) that are unmethylated in normal tissues. DNA methylation levels are visualized as a color gradient heatmap from low (blue) to high (red). The top bar denotes the preservation method while the bottom color bar denotes the tumor types. (b) Correlation coefficients of DNA methylation data between FF and FFPE samples stratified by tumor type. Each colored dot represents the Pearson correlation coefficient between an individual FF-FFPE sample pair. (c) Comparison of DNA methylation levels at the *MLH1* gene locus between FF and FFPE tissues. *Top panel*, genomic tracks of the *MLH1* gene locus, including chromosomal location, location of HM450 DNA methylation probes across the *MLH1* locus and the location of the *MLH1* promoter CpG island. *Lower panel,* heatmap visualization of *MLH1* DNA methylation profiles for FF samples (*left*) and the FFPE samples (*right*). The samples are arranged by tumor type and in the same order for both panels. (d) DNA methylation and gene expression Z-score scatterplots for genes that are epigenetically silenced in both FF and FFPE tissues. The red dots represent the eight genes with differing silencing calls between FF and FFPE samples. The median DNA methylation Z-score is 0.279 while the median expression Z-score is 0.509.

**Figure 6.**
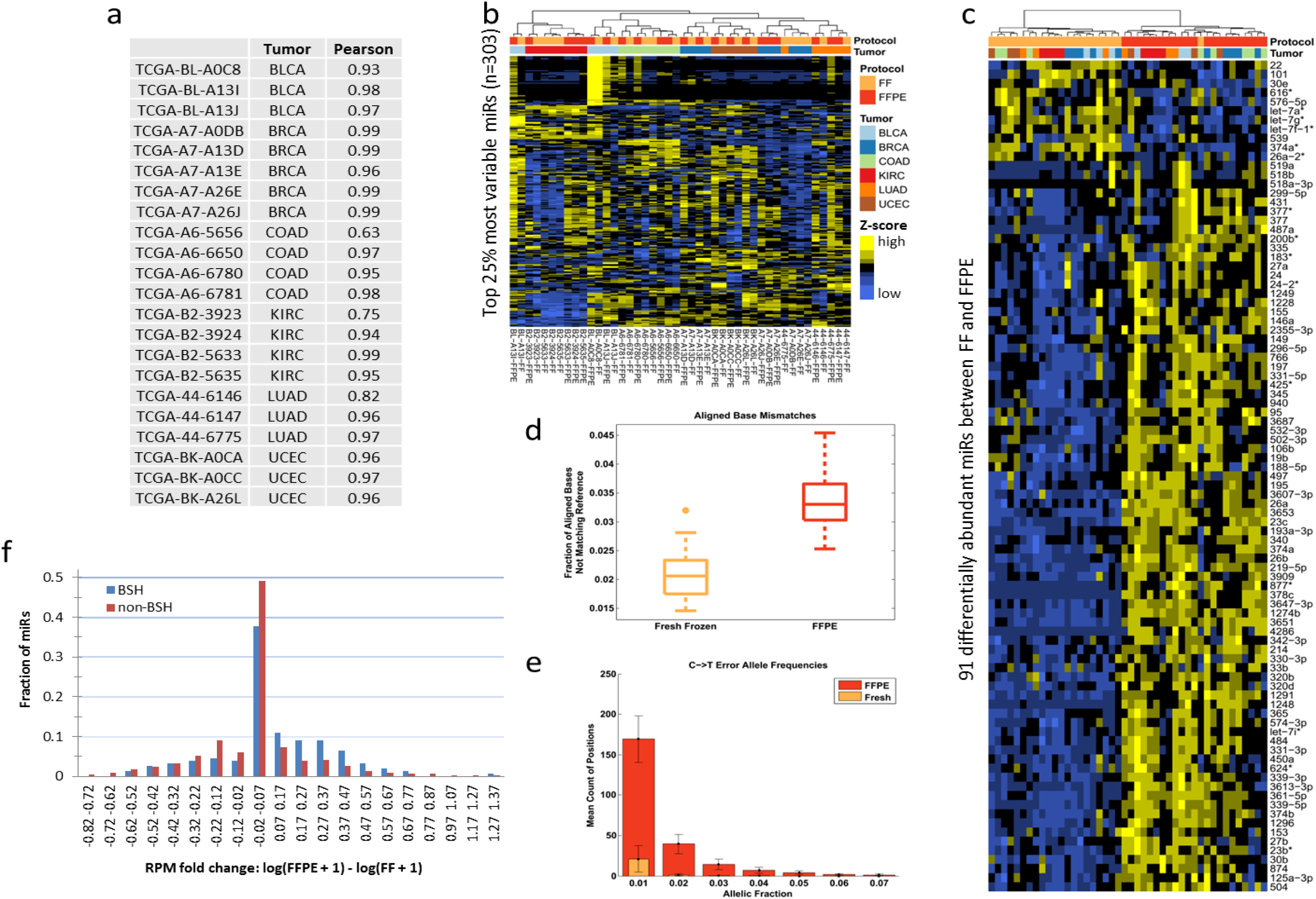
The signature of miRNA expression from FFPE tissues. (a) Pearson correlation between miRNA expression in FF and FFPE sample pairs. (b) Unsupervised hierarchical clustering of all 44 samples using the top 25% most variable miRNAs. (c) Hierarchical clustering using 91 significantly differentially expressed miRs between FF and FFPE. (d) The rate of base errors in aligned miRNA reads as detected by quantifying positions that match the reference. (e) miRNA base mismatches in FFPE are associated with C>T transitions at low allele fractions (<8%). (f) Fold change in miRNA expression from representative patient after classifying miRNA as BSH or non-BSH. Nine of 22 patients exhibited a significant difference between fold changes of non-BSH miRNAs and BSH miRNAs.

To identify factors contributing to DE miRNAs, we investigated the increased base mismatch rate among aligned reads in the FFPE cohort (Figure 6d) with a hypothesis that FFPE-induced base errors, similar to that observed in DNA, may also occur in miRNA. Since miRNA identity is based on unique short sequences, FFPE-induced base modifications may increase miRNA diversity by impairing the accuracy of read mapping to the reference sequence resulting in erroneous mapping of reads to an alternate non-expressed miRNA. In support of this notion, we identified 148 known mature, human miRNAs whose sequence could be converted to that of another miRNA by fewer than four base substitutions (Supplementary Figure 12d; Supplementary Table 5). We refer to these as base substitution homologue (BSH) miRNAs.

While we and others have observed FFPE-induced base errors in RNA (Graw et al., 2015)(Supplementary Figure 13), this is the first report of FFPE-induced base errors in miRNA. Consistent with artifacts found in DNA, we observed an increased number of low allele fraction C>T transitions in miRNA from FFPE samples (Figure 6e). The population of BSH miRNAs detected from FF and FFPE samples (106/148 miRNAs, ≥1 read per million (RPM)) was found to be significantly enriched in the set of miRNAs that are DE between FF and FFPE samples (12/106 BSH miRNA; hypergeometric test; *p*=0.04). However, the total population of DE miRNAs is comprised of a small proportion of BSH miRNA (12/91 DE miRNA) and unsupervised hierarchical clustering after exclusion of the 148 BSH miRNAs did not improve grouping by patient (30/44; Data not shown), suggesting that BSH switching likely does not account for all of the differences we observed between preservation methods. Despite this, nine sample pairs demonstrated a statistically significant increase in expression of BSH compared to Non-BSH miRNA in FFPE (t-test; one-tailed; *p*<=0.05). A representative example from a COAD patient FFPE-FF sample pair (TCGA-A6-5656) is shown (Figure 6f). We do not observe a significant difference in miRNA expression between FF and FFPE for the majority of miRNAs; however, there is a statistically significant increase in the fraction of BSH miRNAs that are expressed at higher levels in the FFPE samples.

## DISCUSSION

MPS has dramatically accelerated the discovery of clinically relevant alterations in FF cancer specimens, and extending this capability to use FFPE requires a thorough understanding of the consequence of formalin-fixation on results. To date, this study represents the most comprehensive comparison of artifacts introduced by FFPE in the context of genomic, transcriptomic and epigenomic alterations in cancer.

This work utilized paired FF and FFPE tumor tissues and blood from 38 patients across six tumor types. We successfully optimized a method for co-isolating DNA and RNA from a single source of scrolls maintaining maximal analyte integrity. Since FFPE-derived nucleic acids are already compromised by formaldehyde, this is a critical first step towards establishing best practices for minimizing additional damage during the extraction.

A central caveat is that the paired FF and FFPE samples were adjacent portions from the same tumor. This spatial separation cofounds subclonal tumor heterogeneity and tumor purity differences with FFPE artifacts. Using LUAD tumors (specifically selected due to their high rate of spatial heterogeneity (Andor et al., 2016)), we were able to quantify the relatively small contribution of spatial heterogeneity for a subset of patients (Supplemental Figure 5). A second caveat of this study is the different nucleic acid extraction methods used for FF and FFPE specimens. This had minimal effect in DNA as evidenced by the high concordance in SCNA and the biology-relevant SMG analysis after appropriate filtering. However, it is unclear if the difference in read distribution in the RNA and the increased miRNA species diversity is a function of FFPE preservation artifacts or the extraction method which could cause selective enrichment of transcripts. Despite this, the hallmarks of FFPE induced artifacts observed in DNA are also observed within the RNA datasets. Specifically, the increase in C>T transitions (both RNA and miRNA) and the increased 3’ bias (RNA degradation) can most likely be attributed to formalin fixation.

Our comparison of FF and FFPE led to several key findings. We successfully used WES data to characterize FFPE preserved tissues; however, it does require optimization of variant calling pipelines. The use of nREF and MCC highlighted how differences in pipelines could alter the detection of variants in FF and FFPE samples. We found that requiring ≥ 20X sequencing coverage or a minimum of 5 variant supporting reads reduced FFPE-specific false positives. Van Allen et al. recently reported a validation rate of 91.5% between mutations found in FFPE and patient matched FF samples (Van Allen et al., 2014), much higher than the ∼30% we observed. This in part, may be due to the pipeline differences a phenomenon we observed in our own cohort (Figure 1c). Ultimately, we found that WES results could successfully be used towards the evaluation of SCNA changes, MSI assessment, and mutational signature analysis without the use of the supporting reads filter.

Assessment of transcriptomic (RNA and miRNA) results from FFPE revealed similar patterns between the two RNA platforms. Overall, there was high correlation in expression profiles between FF and FFPE tissues; however the impact of FFPE was present and we identified RNAs and miRNAs differentially expressed or detected based on preservation status. Similar to DNA, there was evidence of low allele fraction C>T transitions. FFPE resulted in lower quality measures for the total RNA sequencing including reduced reads mapping to coding and UTR regions, a 3’ bias, and smaller insert sizes indicative of degraded RNA. FFPE RNA library preparation does not include steps to fragment nucleic acids; therefore overly degraded RNAs can limit the sequence read length and affect the ability to sequence RNA overall. Despite this, we were able to successfully profile RNA and miRNAs from relatively young FFPE material (1.8 to 3.8 years old) and account for FFPE biases in mRNA results by applying biologically relevant filters to the data.

DNA methylation profiles from FFPE yielded results almost indistinguishable from FF. While assessment of epigenetic silencing did not reveal as strong a correlation between FF and FFPE, we found that variability in RNA expression exerted a stronger influence over epigenetic silencing calls than differences in DNA methylation.

In summary, this work provides readers with access to an optimized FFPE co-isolation protocol, a detailed technical survey of the artifact introduced by FFPE across, genomic, transcriptomic and epigenomic characterization platforms, as well as raw characterization data sets that can be used as a validation cohort to evaluate the performance of individual custom bioinformatic pipelines. This resource is intended to facilitate widespread implementation of best practices for working with FFPE in both clinical and research environments.

## MATERIALS AND METHODS

### Specimens and nucleic acid extraction

Biospecimens were collected at diagnosis from patients with breast carcinoma (BRCA; n=5), kidney renal clear cell carcinoma (KIRC; n=4), colon adenocarcinoma (COAD; n=10), endometrioid adenocarcinomas (UCEC; n=4), lung adenocarcinoma (LUAD; n=12) and urothelial bladder carcinomas (BLCA; n=3) according to consent provided by the relevant institutional review boards. Patients were selected only if their treatment plan required surgical resection, had received no prior treatment for their disease, including chemotherapy or radiotherapy, and if tumor from adjacent portions of the tumor were available as both fresh frozen (FF) and formalin fixed-paraffin embedded (FFPE) specimens. Cases were staged according to the American Joint Committee on Cancer (AJCC) staging system. Each FF and FFPE primary tumor specimen had a companion germline normal specimen that was either blood or blood components (including DNA extracted at the tissue source site). Each FF tumor and germline normal specimen was shipped overnight from one tissue source site using a cryoport that maintained an average temperature of less than -180°C. FFPE tumor specimens were shipped and stored at ambient temperature. FF tumor specimens were embedded in optimal cutting temperature (OCT) medium and a histologic section was obtained for review. Pathologic diagnoses were made at local tissue source sites using diagnostic FFPE sections. Each H&E stained section of frozen OCT embedded tumor processed centrally by TCGA was reviewed by a board - certified pathologist to confirm that the tumor specimen was histologically generally consistent with the diagnosis. Per TCGA protocol requirements, the sections were required to contain at least 60% tumor cell nuclei with less than 20% necrosis for inclusion in the study.

DNA and RNA were co-extracted from FF tumor using a modification of the DNA/RNA AllPrep kit (Qiagen). The flow - through from the Qiagen DNA column was processed using a mirVana miRNA Isolation Kit (Ambion). This latter step generated column purified RNA preparations that included RNA <200 nt suitable for miRNA analysis. The full protocol for DNA and RNA co-extraction from FF tissue is provided as a supplemental document online. DNA and RNA were co-extracted from FFPE tumor using a modification of the Qiagen Allprep FFPE kit. Following deparaffinization, paraffin tissue lysis buffer from the Highpure miRNA kit (Roche) was added to the tissue an incubated on a dry heat block at 55°C for 45 min. The samples were centrifuged to pellet the tissue prior to transfer of the RNA containing supernatant to a fresh tube where the remainder of the Highpure protocol was completed. The remaining tissue was resuspended in Qiagen RLT buffer and the remainder of the Allprep FFPE kit protocol was completed. The full protocol for DNA and RNA co-extraction from FFPE tissue can be found as a supplemental document online. DNA was extracted from blood using either the QIAamp blood midi kit (Qiagen) or by a method selected by the tissue source site.

All analytes were quantified by measuring Abs260 with a UV spectrophotometer (RNA) or by PicoGreen assay (DNA). Analytes were resolved by 1% agarose gel electrophoresis (DNA) or Bioanalyzer RNA6000 nano assay (RNA) to assess analyte integrity. A custom Sequenom SNP panel was utilized to verify tumor DNA and germline DNA were derived from the same patient. Target analyte yield was 6.9 µg of tumor DNA, 5.15 µg RNA, and 4.9 µg of germline DNA. If any yields fell short of these targets, additional extractions were performed and the resulting analytes were pooled until target yield was met or all tissue had been depleted. From FF specimens, DNA analytes with fragmentation resulting in low molecular weight smears or RNA with RIN < 7.0 were excluded in this study. There was no integrity requirement for analytes derived from FFPE specimens.

### Exome sequencing

Exome sequencing was performed on samples from each of the tumor types at one of three Genome Sequencing Centers (GSC; Pipeline 1= Baylor Human Genome Center, Pipeline 2= The Genome Institute at Washington University and Pipeline 3= Broad Institute) using internally developed best practices. The exomes of 38 pairs of FF and FFPE tumor samples, and their respective germline DNA were targeted using HGSC VCRome 2.1 design (42Mb, NimbleGen), Agilent SureSelect All Exon 50Mb or Nimblegen SeqCap v2/v3, and sequenced on Illumina GAIIx or HiSeq 2000/2500 platforms.

### Baylor Human Genome Center

#### Library Preparation

A total of 42 DNA samples (14 triplicate sets consisting of FFPE, fresh frozen (FF), and blood normal DNA) were sequenced at BCM-HGSC. These 14 sets included samples from 10 Colon Adenocarcinoma (COAD) and 4 Kidney Renal Clear Cell Carcinoma (KIRC) patients. Libraries were constructed into Illumina paired-end pre-capture libraries according to the manufacturer’s protocol (*Illumina Multiplexing_SamplePrep_Guide_1005361_D*) with the modifications described below. Libraries were prepared using Beckman robotic workstations (Biomek FX and FXp models). The complete protocol and oligonucleotide sequences are accessible from the HGSC website (https://www.hgsc.bcm.edu/content/protocols-sequencing-library-construction).

Briefly, 0.5 ug of DNA in 70 ul volume was sheared into fragments of approximately 200-300 base pairs with the Covaris E210 system (Covaris, Inc. Woburn, MA) followed by end-repair (NEBNext End-Repair Module; Cat. No. E6050L), A-tailing (NEBNext^®^ dA-Tailing Module; Cat. No. E6053L) and ligation of the Illumina multiplexing PE adaptors with barcode sequences using the ExpressLink™ T4 DNA Ligase (a custom product from Life Technologies). In total, a set of 12 such barcodes were used on these samples. Pre-capture ligation-mediated PCR (LM-PCR) was performed for 6-8 cycles using the Library Amplification Readymix containing KAPA HiFi DNA Polymerase (Kapa Biosystems, Inc., Cat # KK2612). Universal primer IMUX-P1.0 and IMUX-P3.0 were used in the PCR amplification. Purification was performed with Agencourt AMPure XP beads after enzymatic reactions. Following the final XP beads purification, quantification and size distribution of the pre-capture LM-PCR product was determined using the LabChip GX electrophoresis system (PerkinElmer) and gel analysis using AlphaView SA Version 3.4 software.

#### Capture Enrichment

For the hybridization step, four pre-capture libraries were pooled together (∼250 ng/sample, totaling 1 ug per pool). All 14 FFPE samples were co-captured with non-FFPE (FF and normal) samples. The pooled libraries were then hybridized in solution to the HGSC VCRome 2.1 design^1^ (42Mb, NimbleGen) according to the manufacturer’s protocol *NimbleGen SeqCap EZ Exome Library SR User’s Guide (Version 2.2)* with minor revisions. Human COT1 DNA and full-length Illumina adaptor-specific blocking oligonucleotides were added into the hybridization to block repetitive genomic sequences and the adaptor sequences. Post-capture LM-PCR amplification was performed using the Library Amplification Readymix containing KAPA HiFi DNA Polymerase (Kapa Biosystems, Inc., Cat # KK2612) with 12 cycles of amplification. After the final AMPure XP bead purification, quantity and size of the capture library was analyzed using the Caliper LabChip GX electrophoresis system. The efficiency of the capture was evaluated by performing a qPCR-based quality check on the enrichment level of four standard NimbleGen control loci. Successful enrichment of the capture libraries was estimated to range from a 6 to 9 of ΔCt value over the non-enriched samples.

#### Sequencing

Library templates were prepared for sequencing using Illumina’s cBot cluster generation system with TruSeq PE Cluster Generation Kits (Cat. No. PE-401-3001, PE-402-4001) according to the manufacturer’s protocol. Briefly, these libraries were denatured with sodium hydroxide and diluted to 6-9 pM in hybridization buffer in order to achieve a load density of ∼800K clusters/mm^2^. Each library pool was loaded in a single lane of a HiSeq flow cell, and each lane was spiked with 2% phiX control library for run quality control. The sample libraries then underwent bridge amplification to form clonal clusters, followed by hybridization with the sequencing primer. Sequencing runs were performed in paired-end mode using the Illumina HiSeq 2000 and 2500 platforms. Using the TruSeq SBS Kits (Cat. No. FC-401-3001, FC-402- 4001), sequencing-by-synthesis reactions were extended for 101 cycles from each end, with an additional 7 cycles for the index read.

#### Primary Data Processing and Sequencing QC

Initial sequence analysis was performed using the HGSC Mercury analysis pipeline (https://www.hgsc.bcm.edu/content/mercury). In summary, the .bcl files produced on-instrument were first transferred into the HGSC analysis infrastructure by the HiSeq Real-time Analysis module. Mercury then ran the vendor’s primary analysis software (CASAVA) to de-multiplex pooled samples and generate sequence reads and base-call confidence values (qualities), followed by the mapping of reads to the GRCh37 Human reference genome (http://www.ncbi.nlm.nih.gov/projects/genome/assembly/grc/human/) using the Burrows-Wheeler aligner (BWA, http://bio-bwa.sourceforge.net/). The resulting BAM (binary alignment/map) file underwent quality recalibration using GATK (http://www.broadinstitute.org/gatk/), and where necessary the merging of separate sequence-event BAMs into a single sample-level BAM; BAM sorting, duplicate read marking, and realignment to improve in/del discovery all occur at this step.

#### Variant calling pipeline

Primary variant calling was performed on BAM files using Atlas-SNP, Atlas-InDel, and PInDel software. Primary calls for each tumor and normal pair were merged and sequencing depth and variant counts were extracted from each BAM file in the pair for all sites. Primary calls were then filtered for variant quality, coverage, and strand-bias and then split into somatic or germline calls depending on if the variant is called in both tissues or just in the tumor tissue. These filtered calls from each program were merged into the final mutation file (MAF). Copy number aberrations were determined using LOHcate and Varscan2 followed by JISTIC. Further data analysis, statistical comparisons, and plot generation was done using R. The above variants were further characterized across samples and cancer types between the FFPE and FF analytes by performing cross sample variant resolution using Wheeljack.

#### Broad Institute

Exome capture was performed using Agilent SureSelect Human All Exon 50 Mb according to the manufacturers’ instructions. Briefly, 0.5–3 micrograms of DNA from each sample were used to prepare the sequencing library through shearing of the DNA followed by ligation of sequencing adaptors. All whole exome (WES) and whole genome (WGS) sequencing was performed on the Illumina HiSeq platform. Paired-end sequencing (2 x 101 bp for WGS and 2 x 76 bp for WGS) was carried out using HiSeq sequencing instruments; the resulting data was analyzed with the current Illumina pipeline. Basic alignment and sequence QC was done on the Picard and Firehose pipelines at the Broad Institute.

#### The Genome Institute at Washington University

Libraries were prepared using NEB phusion polymerase, indexed and pooled pre- capture. The pools were then captured with Nimblegen Exome v3. UCEC samples had an additional spike-in of probes targeting human papillomavirus (HPV). They were loaded onto lanes of Illumina HiSeq2000 or GAIIx instruments according to the manufacturer’s recommendations. A second round of sequencing was performed to top- up samples that did not meet initial coverage requirements.

#### Non-Reference Variant Calling Enriching for FFPE Artifacts

Exome sequencing data in BAM format for 142 samples from across the 38 cases were downloaded from CGHub [10.1093/database/bau093]. Since these BAMs were generated by 3 different reference-alignment workflows, they were unpacked into FASTQ format using Picard RevertSam v1.123, realigned with BWA-MEM v0.7.10-r789 (runtime arguments: -t 4 -M -R <READGROUP_INFO>) to the GRCh37.p13 reference assembly, and coordinate sorted with samtools sort v1.2. Picard MergeSamFiles v1.123 was used to merge per-readgroup BAMs belonging to the same aliquot, followed by Picard MarkDuplicates v1.123 to tag reads that are likely from the same DNA fragments. Picard HsMetrics v1.123 was used to generate coverage metrics across TCGA GAF4.0b1 genomic intervals [https://tcga-data.nci.nih.gov/docs/GAF/GAF4_0b1].

For calling non-reference variants, BAMs from the same case were pooled together, and the resulting 38 batches were run through samtools mpileup v1.2 (runtime arguments: --no-BAQ --adjust-MQ 50 --min-MQ 1 --min-BQ 20 --ignore-RG --excl-flags UNMAP,SECONDARY,QCFAIL,DUP --output-tags DP,DV,DP4,SP --ext-prob 20 --gap-frac 0.005 --tandem-qual 80 --min-ireads 2 --open-prob 40 --per-sample-mF --platforms illumina) followed by bcftools call v1.2 multi-allelic caller with prior disabled (runtime arguments: --variants-only --multiallelic-caller --prior 0). The runtime arguments selected are more likely to retain FFPE artifacts, at the cost of missing variants at lower allele fractions. On the resulting multi-sample VCFs, additional false-positive filters were applied using bcftools filter v1.3.1. This removed calls seen in the non-TCGA subset of ExAC v0.3.1 [http://exac.broadinstitute.org] with an adjusted minor allele count (AC_Adj) >2, calls supported by lower quality reads (per mpileup metrics: QUAL<10 or (INFO/RPB<0.1 && QUAL<15) or (INFO/MQSB<0.1 && QUAL<15)), calls with depth <10 in at least one of the matched FFPE/Frozen/Blood samples, calls supported by fewer than 2 reads, calls in non-autosomes (X, Y, MT, and unplaced GL* contigs), and calls that are not within a canonical protein-coding exon (per Gencode v19) or <=8bp flanking it. All samples from patient TCGA-A7-A0DB were also removed because it accounted for nearly half of all FFPE-unique events, mostly C>A transversions. Finally bcftools isec v1.3.1 was used to separate the remaining calls as unique to FFPE samples, unique to Fresh Frozen samples, or calls seen in both.

#### Significantly Mutated Gene Analysis

Cross called VCF files were processed using R to extract count, quality, and allele information into an R data.frame which was then analyzed. Variants were filtered based on gene name from the following list of genes previously identified as significantly mutated for these cancer types. ACVR2A, APC, BRAF, CTNNB1, FBXW7, KRAS, PIK3CA, TCF7L2, TP53, ARID1A, BCOR, CHD4, CSMD3, PIK3R1, PTEN, SPOP, AGAP6, EGFR, FSCB, HRNR, KEAP1, KRTAP4, RBM10, STED2, SMARCA4, SPATA31C1, STK11, ZNF679, CDH1, CTCF, KMT2C, MAP3K1, NCOR1, RB1, RPGR, RUNZ1, TBX3, NFE2L2, PBRM1 and VHL. Count, quality, and allele information were plotted using ggplot2.

#### Microsatellite instability (MSI) analyses

Microsatellite instability was evaluated using the Homopolymer Score method and the Mono/Di-nucleotide assay described previously (2012b; Shinbrot et al., 2014).

#### nsNMF Mutational Spectrum Analysis

nsNMF Mutational Spectrum Analysis was performed as described previously (BioRxiv: https://doi.org/10.1101/036541). Briefly, quality filtered MAF files were scored for mutation signatures using an nsNMF mutation solution. Mutation signature scores were then merged with metadata for each sample (sequencing center, cancer type, etc.) and analyzed using R.

#### Heterogeneity analysis

Heterogeneous variants were defined by variant lookup from all available tissues to determine trunk from branch variation. Variants were then grouped and scored using the nsNMF procedure. Signature scores were then merged with metatdata and analyzed using R.

#### Affymetrix SNP 6.0 array and copy number analysis

DNA from each tumor or germline sample was hybridized to Affymetrix SNP 6.0 arrays using protocols at the Genome Analysis Platform of the Broad Institute as previously described (McCarroll et al., 2008). Briefly, from raw .CEL files, Birdseed was used to infer a preliminary copy number at each probe locus (Korn et al., 2008). For each tumor, genome-wide copy number estimates were refined using tangent normalization, in which tumor signal intensities are divided by signal intensities from the linear combination of all normal samples that are most similar to the tumor ((2011) and Tabak B. and Beroukhim R. Manuscript in preparation). This linear combination of normal samples tends to match the noise profile of the tumor better than any set of individual normal samples, thereby reducing the contribution of noise to the final copy-number profile. Individual copy-number estimates then underwent segmentation using Circular Binary Segmentation (Olshen et al., 2004). As part of this process of copy number assessment and segmentation, regions corresponding to germline copy-number alterations were removed by applying filters generated from the TCGA germline samples from the ovarian cancer analysis and from samples of this cohort.

For exome sequencing based copy number analysis segmented copy data was obtained using copy number ratios. These were calculated as the ratio of tumor read depth to the average read depth observed in a panel of normal samples using the tool, RECAPSEG. In the instance where probe level data was used, TCGA Level 2 tangent-normalized probe intensities were used and normalized bait coverage from RECAPSEG were used for the exome sequencing data. GISTIC2.0 was used to map genes to segments for the analysis where gene-level results were presented (Mermel et al., 2011). IGV was used for the visualization of copy number data (Robinson et al., 2011).

#### RNA Sequencing (RNA-Seq)

500 ng of RNA was used as the input for the Illumina TruSeq Total RNA Sample Preparation Kit with Ribo-Zero Gold (RS-122-2301 or RS-122-2302). Libraries were prepped according to manufacturer’s instructions and sequenced two per lane on an Illumina HiSeq2000 machine with a 48x7x48 sequence configuration. All samples were processed as described previously (2012a). Bases and QC assessment of sequencing were generated by CASAVA 1.8. QC-passed reads were aligned to the NCBI build 37 (hg19) human reference genome using MapSplice (Wang et al., 2010) v12_07, and the profile was assessed by Picard Tools v1.64 (http://picard.sourceforge.net/). The aligned reads were translated to transcriptome reference of UCSC hg19 GAF2.1 KnownGenes using UBU v1.0 (https://github.com/mozack/ubu). The abundance of transcripts was then estimated using an Expectation-Maximization algorithm implemented in the software package RSEM (Li and Dewey, 2011) v1.1.13.

RNA-Seq gene quantification data was filtered and processed as described in Zhao et al. (Zhao et al., 2014). The log2 transformed abundance of transcripts in the tumor samples was reported and was used to derive the Pearson correlation between mRNA-Seq and Ribo-Zero-Seq protocols. Hierarchical clustering was performed on the top 3000 most variably expressed genes using centroid linkage and was viewed with Java Treeview v1.1.5r2 (Saldanha, 2004).

To identify fusion transcripts, defuse (v0.6.1, (McPherson et al., 2011)) and chimerascan (v0.4.5, (Maher et al., 2009)) using default parameters, against the human reference genome GRCh37. defuse calls with a probability value of < 0.5 were disregarded. To remove normal transcriptional variants and common alignment artefacts, candidates found in a set of 47 TCGA normal samples processed in the same manner were disregarded. All remaining candidates were annotated using OncoFuse (v1.0.6, (Shugay et al., 2013)) to RefSeq genes. Candidates mapping to regions outside of RefSeq genes were disregarded. Comparison of the Jaccard indices was performed using a paired Mann-Whitney U test.

#### Methylation Studies

We used the Illumina Infinium HumanMethylation450 (HM450) platform to obtain DNA methylation profiles for TCGA DNA samples derived from fresh-frozen (FF) and formalin-fixed, paraffin embedded (FFPE) tissues. The HM450 assay analyzes the DNA methylation status of 482,421 CpG dinucleotides and 3,091 non-CpG sites. The platform also contains 65 probes assigned to known single nucleotide polymorphisms (SNPs). Collectively, the HM450 platform covers 99% of NCBI RefSeq genes including promoter, 5’ and 3’ gene body regions, as well as CpG sites located outside of gene coding regions. Probes are located within 80% of core promoters (within 200 bp of the transcription start site (TSS)), 94% of distal promoters (greater than 1,500 bp from the TSS) in the human genome) and 97% of all gene bodies. The HM450 assay contains probes to measure DNA methylation of CpG islands, CpG island shores and non-CpG island regions in the human genome. CpG island regions are annotated as regions greater than 500 bases, with a G:C content >50% and a CpG observed/expected ratio of ≥ 0.6. CpG island shores are characterized as 0-2 kb regions both upstream (“north”) and downstream (“south”) of CpG islands, and CpG island shelves are the 0-2 kb regions both upstream and downstream of CpG island shores. A total of 26,658 CpG islands (94%) are interrogated on the HM450 array, together with 26,249 (95%) north CpG island shores, 25,761 (93%) south CpG island shores, 23,965 (86%) north CpG island shelves and 24,018 (87%) south CpG island shores.

Analytes were prepared for HM450 hybridization as follows. Genomic DNAs from FFPE tissues are treated with sodium bisulfite. After bisulfite conversion, each sample is repaired using the Illumina Restoration Solution as recommended by the manufacturer. The entire restored sample is then whole genome amplified (WGA) and enzymatically fragmented. Samples are then hybridized overnight to a 12 sample BeadChip, in which the WGA-DNA molecules anneal to locus-specific DNA oligomers linked to individual bead types. The oligomer probe designs follow the Infinium I and II chemistries, in which base extension follows hybridization to a locus-specific oligomer. With respect to the Infinium I probes, there are two different bead types for each locus, one with an oligomer that anneals specifically to the methylated version of the locus, while the other oligomer anneals to the unmethylated version of the locus. The probes terminate complementary to the interrogated CpG site for methylated loci, or complementary to the TpG for unmethylated alleles. A matched oligomer-template DNA molecule hybrid will allow for the incorporation of a labeled nucleotide immediately upstream (5’) to the interrogated CpG (or TpG) site. However, if the probe and template are mismatched, then primer extension will not occur. Adenine and thymine nucleotides are labeled with cy5 (red), while cytosine nucleotides are labeled with cy3 (green). No insertion of guanine nucleotides occurs in Infinium I assays. Of note, the identity of the dye is representative of the nucleotide adjacent to the CpG dinucleotide. The methylation discrimination is derived from separate measurements from the two different types of beads present for each locus. For some loci, both measurements will be cy3, and for others both will be cy5. The Infinium type II chemistry is a true two-color system. A matched oligomer-template DNA molecule hybrid will allow for the incorporation of a labeled nucleotide at the interrogated C or T of the CpG site. Adenine nucleotides labeled with cy5 (red) are incorporated across from unmethylated (TpG) sites, while guanine nucleotides labeled with cy3 (green) are incorporated across from methylated (CpG) sites.

BeadArrays were scanned and the raw signal intensities were extracted from the *.IDAT files using the R package *methylumi*. The intensities were corrected for background fluorescence and red-green dye-bias using the methods described previously. (Triche et al., 2013). The beta value was calculated as (M/(M+U)), in which M and U refer to the (pre-processed) mean methylated and unmethylated probe signal intensities, respectively. Measurements in which the fluorescent intensity was not statistically significantly above background signal (detection p value > 0.01) were removed from the data set. Since the HM450 assay utilizes a hybridization-based strategy, it is important to ascertain the specific CpG dinucleotides that are associated with known SNPs, as one could analyze the SNP rather than the DNA methylation at the CpG dinucleotide, and would then lead to misrepresentation of the data and conclusions. Therefore, probes that overlap with known SNPs as well as repetitive elements were masked prior to data analyses. Specifically, we masked all HM450 probes that have common SNPs with a minor allele frequency of greater than 1% (UCSC criteria) at the targeted CpG site, as well as probes with common SNPs (MAF>1%) within 10 bp of the targeted CpG site. With regards to repetitive elements, HM450 probes that were within 15 bases of the CpG lying entirely within a repeat region were masked prior to data analyses. Generally, high-quality samples yield in excess of 99% successful measurements.

#### Unsupervised Clustering Analysis of Methylation Results

We first selected CpG loci that are unmethylated (median beta value <= 0.2) in normal tissues. We previously identified a set of 12,862 CpG probes that were unmethylated in normal tissues across the 12 tissue types analyzed for the TCGA Pan-Cancer project. The 12 tissue types included in the TCGA Pan-Cancer analysis include all six tissue types used in this report. We used this set of 12,862 CpG probes to perform two-dimensional unsupervised clustering of the paired FF and FFPE across all six tissue types. As these loci are mostly within CpG islands that remain constitutively unmethylated in normal tissues, we dichotomized the data using a beta value threshold of 0.3, in which tumors with a beta value ≥ 0.3 are designated as methylated, while tumors with a beta value <0.3 are designated as unmethylated. The dichotomization greatly ameliorates the effect of tumor sample purity on the clustering and removes a substantial portion of residual batch/platform effects that are mostly reflected in small variations near the two extremes of DNA methylation (0 and 1). We performed two-dimensional unsupervised hierarchical clustering using Ward’s method on the Jaccard Distance, a distance measure that best suits binary data. This was visualized using a heatmap. We annotated the heatmap with the sample preservation method (FF or FFPE), tissue type and FF-FFPE pair identifiers.

#### Epigenetic Silencing Calling from Methylation and RNA Seq Results

We downloaded RSEM Normalized Level 3 RNA-seq data for 32 FF-FFPE pairs from the TCGA Data Portal. The RNA-seq data were generated on the Illumina HiSeq platform, mapped with the RSEM algorithm and normalized so that the third quartile for each sample equals 1000. Entrez gene IDs were used for mapping to genomic locations using the R/Bioconductor *Homo.sapiens* package, and DNA methylation probes that mapped to [-1500, +200] of the TSS for different transcripts for each gene were studied for the corresponding gene. We only considered the 12,862 CpG probes that are unmethylated in normal tissues, and were able to map 12,508 CpG probes to the aforementioned genomic range for a known transcript. For each probe/gene pair, the tumors were first divided into unmethylated/methylated groups with a beta value range of (0,0.2) and (0.3,1) respectively. We then log_2_-transformed the expression data [log_2_ (RSEM+1)], and computed the mean (X_u_) and standard deviation (S_u_) expression levels of the corresponding gene for the unmethylated group. Then, we computed the mean (X_m_) expression of the methylated group and normalized it by N = (X_m_ – X_u_) / S_u_, in which *N* indicates the number of standard deviations by which the mean expression of the methylated group is away from that of the unmethylated group. We then computed an average N for each gene. Only genes for which the methylated group had an averaged expression of ≥2 standard deviations lower than the unmethylated group were called epigenetically silenced.

For each tumor sample, we next considered all probes where correlation with expression was detected, and then required the sample to show DNA methylation (beta value > 0.3) in more than half the probes to be called silenced for each specific gene. We called a gene as epigenetically silenced across all samples only if the gene was silenced in at least four samples and the highest beta value was at ≥0.7 across the sample set. We performed this analysis separately on data for FF and FFPE samples.

In order to compare the influence of DNA methylation and expression on the epigenetic silencing calls, we calculated a Z-score using the following method: For each gene we calculated the standard deviation (S_E_) of gene expression across all 32 FF-FFPE pairs (64 total samples). Next, we calculated the mean (A_E_) of the absolute pair-wise difference in expression for each gene between the FF-FFPE samples. We calculated the Z-score as Z = A_E_ / S_E_. We repeated the process for the DNA methylation values and then generated a scatterplot of the Expression Z-scores (y-axis) versus the DNA Methylation Z-scores (x-axis). Scatterplots were generated for both the set of epigenetically silenced genes in common between FF and FFPE samples, as well as for the set of genes that were not in agreement between FF and FFPE samples.

#### miRNA Sequencing (miRNA-Seq)

##### Library construction and sequencing

Two micrograms of total RNA per sample were arrayed into 96-well plates, with controls as described below. RNA entering library construction was required to have at least a minimum quality on the BCR submission documentation. miRNA-Seq libraries were constructed using a plate-based protocol developed at the British Columbia Genome Sciences Centre (BCGSC). Negative controls were added at three stages: elution buffer was added to one well when the total RNA was loaded onto the plate, water to another well just before ligating the 3’ adapter, and PCR brew mix to a final well just before PCR. A 3’ adapter was ligated using a truncated T4 RNA ligase2 (NEB Canada, cat. M0242L) with an incubation of 1 hour at 22°C. This adapter is adenylated, single-strand DNA with the sequence 5’ /5rApp/ ATCTCGTATGCCGTCTTCTGCTTGT /3ddC/, which selectively ligates to miRNAs. An RNA 5’ adapter was then added, using a T4 RNA ligase (Ambion USA, cat. AM2141) and ATP, and was incubated at 37°C for 1 hour. The sequence of the single strand RNA adapter is 5’- GUUCAGAGUUCUACAGUCCGACGAUCUGGUCAA-3’.

When ligation was complete, 1^st^ strand cDNA was synthesized using Superscript II Reverse Transcriptase (Invitrogen, cat.18064 014) and RT primer (5’-CAAGCAGAAGACGGCATACGAGAT-3’). This is the template for the final library PCR, into which we introduce index sequences to enable libraries to be identified from a sequenced pool that contains multiple libraries. Briefly, a PCR brew mix was made with the 3’ PCR primer (5’-CAAGCAGAAGACGGCATACGAGAT-3’), Phusion Hot Start High Fidelity DNA polymerase (NEB Canada, cat. F-540L), buffer, dNTPs and DMSO. The mix was distributed evenly into a new 96-well plate. A Biomek FX robotic platform (Beckman Coulter, USA) was used to transfer the PCR template (1^st^ strand cDNA) and indexed 5’ PCR primers into the brew mix plate. Each indexed 5’ PCR primer, 5’-AATGATACGGCGACCACCGACAGNNNNNNGTTCAGAGTTCTACAGTCCGA-3’, contains a unique six-nucleotide ‘index’ (shown here as N’s), and is added to each well of the 96-well PCR brew plate. PCR was run at 98°C for 30 sec, followed by 15 cycles of 98°C for 15 sec, 62°C for 30 sec and 72°C for 15 sec, and finally a 5 min incubation at 72°C. Library quality was then checked across the whole plate using a Caliper LabChipGX DNA chip. PCR products were pooled (16 libraries per pool), then were size selected to remove larger cDNA fragments and smaller adapter contaminants, using a 96-channel automated size selection robot that was developed at the BCGSC. After size selection, each pool was ethanol precipitated, quality checked using an Agilent Bioanalyzer DNA1000 chip and quantified using a Qubit fluorometer (Invitrogen, cat. Q32854). Each pool was then diluted to a target concentration for cluster generation and loaded into a single lane of a HiSeq 2000 flow cell. Clusters were generated, and lanes were sequenced with a 31-bp main read for the insert and a 7-bp read for the index.

##### Preprocessing, alignment and annotation

Briefly, the sequence data were separated into individual samples based on the index read sequences, and the reads underwent an initial QC assessment. Adapter sequence was trimmed off, and the trimmed reads for each sample were aligned to the NCBI GRCh37-lite reference genome. Below we describe these steps in more detail.

Routine QC assesses a subset of raw sequences from each pooled lane for the abundance of reads from each indexed sample in the pool, the proportion of reads that possibly originate from adapter dimers (i.e. a 5’ adapter joined to a 3’ adapter with no intervening biological sequence) and for the proportion of reads that map to human miRNAs. Sequencing error was estimated by a method originally developed for SAGE (Khattra et al., 2007).

Libraries that pass this QC stage were preprocessed for alignment. While the size-selected miRNAs vary somewhat in length, typically they are ∼21 bp long, and so are shorter than the 31-bp read length. Given this, each sequence read extends some distance into the 3’ sequencing adapter. Because this non-biological sequence can interfere with aligning the read to the reference genome, 3’ adapter sequence was identified and removed (trimmed) from a read. The adapter-trimming algorithm identified as long an adapter sequence as possible, allowing a number of mismatches that depends on the adapter length found. A typical sequencing run yields several million reads; using only the first (5’) 15 bases of the 3’ adapter in trimming makes processing efficient, while minimizing the chance that a miRNA read will match the adapter sequence.

The algorithm first determines whether a read sequence should be discarded as an adapter dimer by checking whether the 3’ adapter sequence occurs at the start of the read. For reads passing this stage, the algorithm then attempts to identify an exact 15-bp match anywhere within the read sequence. If it cannot, it then retries, starting from the 3’ end, and allowing up to 2 mismatches. If the full 15bp is not found, decreasing lengths of adapter are checked, down to the first 8 bases, allowing one mismatch. If a match is still not found, from 7 bases down to 1 base is checked, with an exact match required. Finally, the algorithm will trim 1 base off the 3’ end of a read if it happens to match the first base of the adapter. This is based on two considerations. First, it is preferable to get a perfect alignment than an alignment that has a potential one-base mismatch. Second, if only 1 base of adapter was found in the sequence read, the read is likely too long to be from a miRNA and the effect of the trimming on its alignment would not affect this sample’s overall miRNA profiling result.

After each read has been processed, a summary report was generated containing the number of reads at each read length. Because the shortest mature miRNA in miRBase v16 is 15 bp, any trimmed read that was shorter than 15bp was discarded; remaining reads were submitted for alignment to the reference genome.

BWA (Li and Durbin, 2009) alignment(s) for each read were checked with a series of three filters. A read with more than 3 alignments was discarded as too ambiguous. For TCGA quantification reports, only perfect alignments with no mismatches were used. Based on comparing expression profiles of test libraries (data not shown), reads that fail the Illumina basecalling chastity filter were retained, while reads that have soft-clipped CIGAR strings were discarded.

For reads retained after filtering, each coordinate for each read alignment was annotated using the reference databases (Supplemental Table 6), and required a minimum 3-bp overlap between the alignment and an annotation. In annotating reads we addressed two potential issues. First, a single read alignment can overlap feature annotations of different types; second, a read can have up to three alignment locations, and each alignment location can overlap a different type of feature annotation. By considering heuristically determined priorities (Supplemental Table 6), we resolved the first issue by giving each alignment a single annotation. We resolved the second by collapsing multiple annotations to a single annotation, as follows.

If a read has more than one alignment location, and the annotations for these are different, we use the priorities from Supplemental Table 6 to assign a single annotation to the read, as long as only one alignment is to a miRNA. When there are multiple alignments to different miRNAs, the read is flagged as cross-mapped (de Hoon et al., 2010), and all of its miRNA annotations are preserved, while all of its non-miRNA annotations are discarded. This ensures that all annotation information about ambiguously mapped miRNAs is retained, and allows annotation ambiguity to be addressed in downstream analyses. Note that we consider miRNAs to be cross-mapped only if they map to different miRNAs, not to functionally identical miRNAs that are expressed from different locations in the genome. Such cases are indicated by miRNA miRBase names, which can have up to 4 separate sections separated by "-", e.g. hsa-mir-26a-1. A difference in the final (e.g. ‘-1’) suffix denotes functionally equivalent miRNAs expressed from different regions of the genome, and we consider only the first 3 sections (e.g. ‘hsa-mir-26a’) when comparing names. As long as a read maps to multiple miRNAs for which the first 3 sections of the name are identical (e.g. hsa-mir-26a-1 and hsa-mir-26a-2), it is treated as if it maps to only one miRNA, and is not flagged as cross-mapped.

From the profiling results for a tumor type, for a minimum of approximately 100 samples, we identify the depth of sequencing required to detect the miRNAs that are expressed in a sample by considering a graph of the number of miRNAs detected in a sample as a function of the number of reads aligned to miRNAs. For the current work, a library from a sequenced pool was required to have at least 750,000 reads mapped to miRBase annotations. For any sequencing run that fails to meet this threshold, we sequence the sample again to achieve at least the minimum number of miRNA-aligned reads.

Finally, for each sample, the reads that correspond to particular miRNAs are summed and normalized to a million miRNA-aligned reads to generate the quantification files that are submitted to the DCC. Quantification files include information on variable 5’ and 3’ read alignment locations, which can reflect isoforms, adapter trimming and RNA degradation.

#### Notes for implementation

Our adapter trimming code is available at: http://www.bcgsc.ca/platform/bioinfo/software/adapter-trimming-for-small-rna-sequencing. For alignment, we use BWA 0.5.7: bwa samse –n 10. .bam files must be converted to .sam format to be used in the annotation pipeline: $samtools view -h $bam > $sam. The current release of the annotation pipeline is available at: http://www.bcgsc.ca/platform/bioinfo/software/mirna-profiling. This code uses a .sam file containing BWA alignments of trimmed reads for a sample as input.

There are two configuration files which point the code to all of the necessary inputs:

- **db_connections.cfg**: contains parameters to access MySQL databases containing the necessary UCSC and miRBase information. The db_name field is used when providing the database source for various script parameters. You must have a database connection to a miRBase instance and a UCSC database instance for annotations of miRNAs and other non-coding RNAs respectively. In db_connections.cfg, the server name of the database is the <HOST> parameter, and the login and password are the <USER> and <PASSWORD> parameters.
- **profile.sh**: points to the appropriate perl to use for the scripts. Run source on this to generate the environment. e.g.source $BASEDIR/v0.2.7/config/profile.sh

#### Other requirements

Perl (perl-5.10-x86_64), samtools (0.1.7) and R (R-2.12.0) must be available on your system. In addition, perl requires the MySQL DBI library.

#### Differential miRNA expression and clustering analyses

To identify miRNAs that are differentially abundant between fresh-frozen and FFPE, we ran a two-class paired analysis using SAMseq, with a read-count input matrix and an FDR threshold of 0.05. We filtered the results to retain only those with a SAMseq qvalue of 0 and then removed miRs with median expression less than 1 RPM, or with Wilcoxon adjusted p-value greater than 0.1. When clustered, these 91 differentially abundant miRNAs segregate the FF from FFPE samples with the exception of sample A6-5656 which has the lowest Pearson correlation with between FF and FFPE.

To identify samples that preferentially group together we used hierarchical clustering with pheatmap v1.0.2 in R. The input was a reads-per-million (RPM) data matrix for the 303 miRBase v16 5p or 3p mature strands that had the largest variances across the cohort. We transformed each row of the matrix by log_10_(RPM + 1), then used pheatmap to scale the rows. We used Ward.D2 for the clustering method with correlation and Euclidean as the distance measures for clustering the columns and rows respectively.

To investigate the possibility that the greater species diversity seen in FFPE miRNA samples may be an artifact caused by identifying degraded mRNA as miRNA species, sequences for miRNAs that were differentially expressed between the fresh-frozen and FFPE samples were checked for genomic CDS multi-mapping. The expectation would be that miRNAs in FFPE samples with significantly higher levels of expression may have derived from degraded mRNA coding sequence. All differentially expressed miRNAs were mapped to the human CDS (ensemble) using BLAT. None of the 43 over-expressed miRNAs obtained complete-sequence matches to the human CDS indicating that the differences in expression between fresh-frozen and FFPE samples were not a product of mRNA degradation.

To identify the base substitution homologue (BSH) miRNAs, we first exhaustively aligned the sequences of all miRNA sequences listed in miRBase 16 against each other using ggsearch36 from the Fasta 3 package (Pearson, 2000) with the following settings: -d 1 show one best alignment;-C 1000 length of name abbreviation in alignments; -m 8 tabular output.

#### AVAILABILITY

Data for analysis is available in the GDC legacy archive repository (https://portal.gdc.cancer.gov/legacy-archive/)

## Supporting information

TCGA Research Network List

Supplemental Table 1 - BCR Extraction Summary

Supplemental Table 3 - List of Fusions

Supplemental Table 2 - Concordance Summary

Supplemental Table 4 - Differentially expressed miRNA

Supplemental Table 5 - Edit homologue miRs

Supplemental Table 6 - miRNA Annotation Priorities

## ACKNOWLEDGMENTS

Acknowledgements: We are grateful to all patients who contributed to this study and to I. Felau and M. Sheth for administrative support.

## AUTHOR CONTRIBUTIONS

Designed and supervised project: RT, EJZ RNA and DNA isolation: EJZ, JGF

DNA sequencing: CM, MM, DAW, RAG, DM, DM, JH, CS, ES, NK, LX, BF

RNA sequencing: KAH, WZ

miRNA sequencing and analysis: RB, RDC, DB, AJM, AC

SNP6.0 assay: ADC, CS, BM

DNA methylation assay: PWL, DJW

DNA mutation calling: HVD, KRC, CK, CM, MM, CS

DNA mutation analysis: KRC, CK, CN, JSR, HVD

Copy number analysis: BM, AC, ML, PS

Fusion analysis: CKYN RNA analysis: KAH, WZ

DNA methylation analysis: PWL, DJQ, MSB, TH

Provided program oversight: CH, JCZ, RT

Provided critical input into the project: RA, CC, GG, PS, NS, JSR

Wrote the manuscript: EJZ, KAH, PWL, DJW, ADC, HVD, DAW, CC, AJM, RDC, RB

All authors reviewed the manuscript

