## Supplemental Materials for "Comprehensive characterization of genomic, transcriptomic and epigenomic artifacts introduced in formalin-fixed, paraffin-embedded tissues"

**Supplemental Text Figures and Tables**

**Supplemental figure 1** Extraction method development and analyte quality control results. (a) Representative nucleic acid integrity results from high quality Thymus control tissues. Top- DNA integrity was visualized following electrophoresis on a 1% agarose gel stained with ethidium bromide. Bottom- RNA integrity as assessed using Agilent Bioanalyzer Nano6000 chips. RIN values were monitored from FF while %DV200 was used to evaluate integrity of FFPE. All samples shown are FFPE unless otherwise noted. (b) Overview of FF and FFPE co-extraction methods used in this study. (c) Summary of FFPE block characteristics and nucleic acid extraction results. (d) Cross platform distribution of paired FF and FFPE analytes.

**Supplemental figure 2** The mutational signature of FFPE as viewed through a minimally filtered variant calling pipeline. Distribution of single base substitutions (SBS) by tissue type, base, VAF, supporting reads and sequencing depth from non-reference base calling. SBS unique to FFPE (Purple) and shared between FF and FFPE (Yellow) are shown.

**Supplemental figure 3** Detecting MSI in FFPE tumors. (a) Homopolymer analysis of FFPE vs Frozen samples. The homopolymer score for each tumor is plotted on the y axis in a stacked bar graph (see methods) while each individual sample, grouped by tumor type, is on the X axis. MSI designations assigned by TCGA using the Mono/Di-Nucleotide PCR based assay are labeled on the top of the graph. Abbreviations: S, microsatellite stable; L, microsatellite low; H, microsatellite high, colorectal cancer (COAD) and endometrial carcinoma (UCEC).

**Supplemental figure 4** nsNMF Mutational Signature analysis in FFPE. (a) Heatmap of FF signatures with dominant penetrance across cancer types. The ribbon represents disease. Shown at right are biological contexts associated with the individual signatures as determined through literature survey. (b) Correlation coefficients between FF and FFPE samples across the different pipelines for each signature. Max p-value:< 8e - 2 observed in the MCC call set for signature 1. All other groups are nominally significant.

**Supplemental figure 5** Signature of spatial heterogeneity from frozen lung adenocarcinoma (LUAD) tumors. (a) The spectrum of SBS that were found unique to FFPE (top) or unique across adjacent frozen portions (bottom) from seven LUAD patients. Shown are variants with a VAF <20%, plotted by trinucleotide sequence context. (b) Full nsNMF heatmap ordered by VAF (green, All; blue, >20% VAF; purple, <20% VAF) and then specimen association (red, FFPE unique; orange, shared across frozen; yellow, unique across frozen). (c) nsNMF heatmap as shown in (b), only expanded to show results from <20% VAF. (d) Left- Quantitation of signature 5 penetrance by sample group across the VAF bins. Right- Quantitation of signature 6 penetrance by sample group across the VAF bins.

**Supplemental figure 6** Probe level detail for SCNAs quantified from SNP6, WES and WGS analyses. Probe level data (Left) and copy number segments from probe data (Right) from two representative patients (a) TCGA-A6-3810 and (b) TCGA-A7-A26J are shown. Platform specific probes are defined as follows: SNP6 probes are SNP and CN probes on the array, WES probes are exome capture probes, and WGS "probes" are computationally derived from evenly spaced 1 kb bins.

**Supplemental figure 7** QC metrics from mRNA sequencing. (a) Percent of reads mapped to exonic, intronic, intergenic and unaligned regions. (b) Median coefficient of variation (CV) coverage. (c)

Uniformity of coverage across normalized transcript length. (D) Median 5' to 3' coverage bias. (e) Median insert size on average (left) and by individual tumor type (right).

**Supplemental figure 8** mRNA sequencing of FF and FFPE specimens. (a) Pairwise correlation of transcript abundance from individual patient samples. Principle component analysis by (b) preservation method and (c) tissue type.

**Supplemental Figure 9** Biology-specific gene filters show reduced FFPE effects on RNA transcriptome concordance within sample pairs. (a) Principal component plots of all expressed genes or using biology-specific gene lists from the previously published TCGA marker papers for each tumor type. Paired T-tests of the PC1 loadings between FF and FFPE and the median of the differences were calculated. Significant p-values indicate FFPE effects on gene expression values, non-significant p-values indicate limited or no effect of FFPE on gene expression. (b) Using the same approach as in (a), except all samples combined in principle component plots of all variably expressed genes, variably expressed genes after subtraction of the FFPE-associated genes, variably expressed genes within only FF, and variably expressed genes with only FFPE.

**Supplemental figure 10** Density distribution and differential DNA methylation between paired FF and FFPE samples. (a) Density distribution of the probe-wise DNA methylation beta value differential between each FF-FFPE sample pair. The vertical dashed line is the median FF-FFPE differential (-0.0168). (b) Scatterplot of differential DNA methylation between FF and FFPE samples as a function of HM450 probe CpG density. The scatterplot displays DNA methylation variation between FF and FFPE samples as a function of HM450 probe Observed/Expected CpG density. Observed/Expected CpG density is calculated as the ratio of the observed number of CpGs to the expected number of CpGs multiplied by the length of the probe (50bp). A negative value on the y-axis represents higher DNA methylation in FFPE samples, whereas a positive value represents higher DNA methylation in FF samples. The loess smoothing line in red shows the variation of the beta value DNA methylation difference between FF and FFPE samples as a function of probe CpG density. The blue horizontal dashed line indicates no difference in DNA methylation between FF and FFPE tissues.

**Supplemental figure 11** Cross-platform integrative analysis using FFPE specimens. DNA methylation and RNA expression levels detected from three representative genes *ZNF549* (left), *RNLS* (middle) and *CHFR* (right). For each gene, unmethylated samples are shown as black closed circles regardless of preservation method. Patients that demonstrated DNA methylation in the target gene are shown in color where open circles representing FF data points and closed circles represent FFPE. FF/FFPE pairs are connected by solid lines. DNA methylation density is plotted on the x-axis while RNA log2(RSEM+1) expression values are plotted on the y-axis.

**Supplemental figure 12** The signature of miRNA expression from FFPE samples. (a) Total reads vs. miRNA aligned reads. (b) miRNA species detected vs. number of miRNA aligned reads. (c) Hierarchical clustering after removing the differentially abundant miRNAs in (Fig. 6b) from (Fig. 6a). (d) Identification of 'edit distance' miRNA using pairwise comparison of similar reference miRNAs. Shown are miRNAs that were less than 3 base substitutions different from one another.

**Supplemental figure 13** Quantifying base errors in mRNA sequencing results from FFPE specimens. Limiting to positions within the transcriptome, we determined the number of C>T base changes relative to the reference. Allelic fraction of C>T changes at each locus were calculated and the mean was determined across FF and FFPE samples per variant allele frequency bin.

**Supplemental table 1** Summary of nucleic acid extraction results.

**Supplemental table 2** WES Summary of SBS and INDEL concordance between FF and FFPE.

**Supplemental table 3** List of all gene fusions detected through RNA sequencing.

**Supplemental table 4** List of differentially expressed miRNA.

**Supplemental table 5** Summary of edit distance between BSH miRNA transcripts.

**Supplemental table 6** miRNA annotation priorities.

### Supplementary Figure 1

**a**

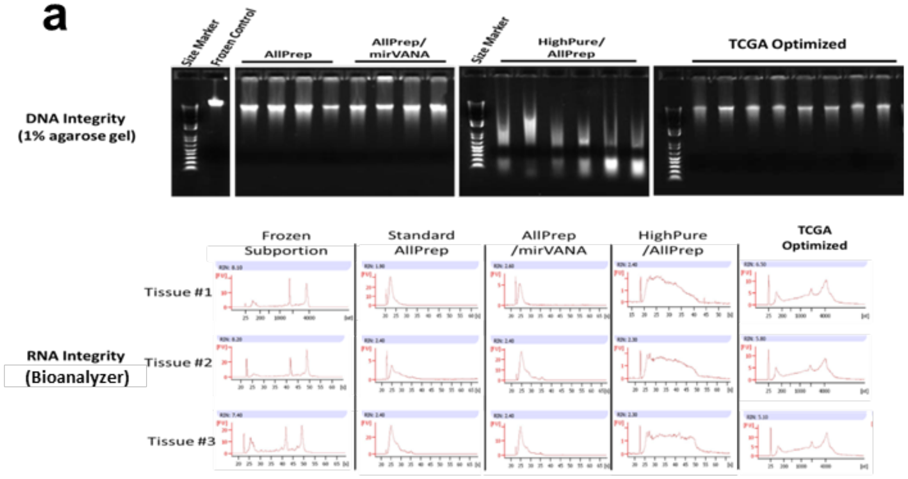


**b**

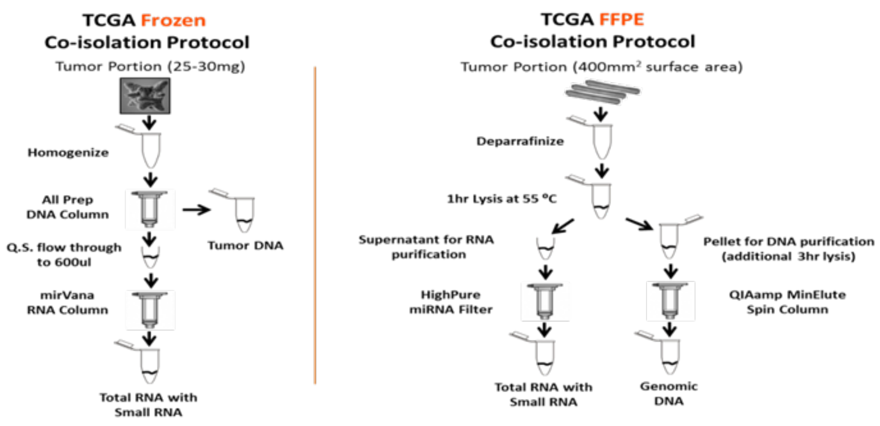


**c**

| Tumor Type | Number of Patients | Tissue Time in 10% Formalin (minutes) | Age of FFPE Tissue Block (years) | % Tumor Nuclei | % Necrosis | Number of Pooled Extractions | Tumor Surface Area Pooled (mm2) | Tumor Surface Area Spanned in Analyte Pool (mm2) | DNA Yield / Extraction (ug per (400mm2) | RNA Yield / Extraction (ug per 400mm2) | RNA Integrity (RIN) | RNA Integrity (DV200) |
|---|---|---|---|---|---|---|---|---|---|---|---|---|
| Colon Adenocarcinoma | 10 | 961.7 +/- 636.54 | 2.86 +/- 0.72 | 74.46 +/- 11.15 | 4.82 +/- 3.89 | 3.2 +/- 1.03 | 1280 +/- 413.12 | 1280 +/- 413.12 | 4.67 +/- 2.17 | 12.03 +/- 7.62 | 2.36 +/- 0.18 | 55 +/- 9.35 |
| Endometrial Carcinoma | 4 | 703.5 +/- 651.56 | 2.64 +/- 0.45 | 71.53 +/- 8.5 | 2.8 +/- 4.1 | 3.5 +/- 1.91 | 1400 +/- 765.94 | 1600 +/- 1131.37 | 4.19 +/- 2.22 | 13.91 +/- 6.34 | 2.43 +/- 0.17 | 55.75 +/- 12.61 |
| Lung Adenocarcinoma | 12 | 780.25 +/- 562.32 | 2.97 +/- 0.63 | 72.64 +/- 6.73 | 5.36 +/- 5.69 | 3.17 +/- 1.34 | 1266.67 +/- 534.85 | 1400 +/- 692.82 | 3.78 +/- 1.69 | 11.68 +/- 5.7 | 2.42 +/- 0.06 | 51.58 +/- 9.1 |
| Bladder Urothelial Carcinoma | 3 | 432.33 +/- 170.2 | 2.72 +/- 0.24 | 89.18 +/- 5.46 | 2.49 +/- 2.49 | 4 +/- 0 | 1600 +/- 0 | 1600 +/- 0 | 3.75 +/- 0.95 | 13.01 +/- 3.59 | 2.33 +/- 0.06 | 41.67 +/- 1.53 |
| Kidney Renal Clear Cell Carcinoma | 4 | 437 +/- 150.1 | 2.89 +/- 0.12 | 89.86 +/- 6.8 | 0.83 +/- 1.18 | 5.5 +/- 3 | 2200 +/- 1200 | 2200 +/- 1200 | 2.63 +/- 1.46 | 6.12 +/- 2.29 | 1.9 +/- 0.54 | 40.25 +/- 9.54 |
| Breast Invasive Carcinoma | 5 | 480.8 +/- 144.8 | 2.66 +/- 0.5 | 74.16 +/- 5.02 | 4.03 +/- 6.33 | 4 +/- 2 | 1600 +/- 800 | 1600 +/- 800 | 1.66 +/- 1.6 | 4.21 +/- 2.44 | 2.32 +/- 0.24 | 45.6 +/- 5.18 |
| **Total/Average** | **38** | **716.92** | **2.84** | **76.32** | **4.07** | **3.63** | **1452.63** | **1515.79** | **3.65** | **10.54** | **2.33** | **50.16** |

**d**

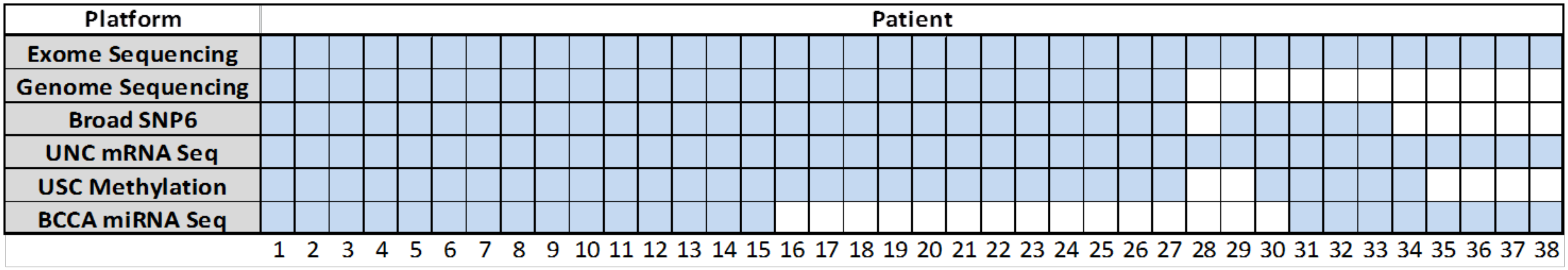

**Supplementary Figure 2**

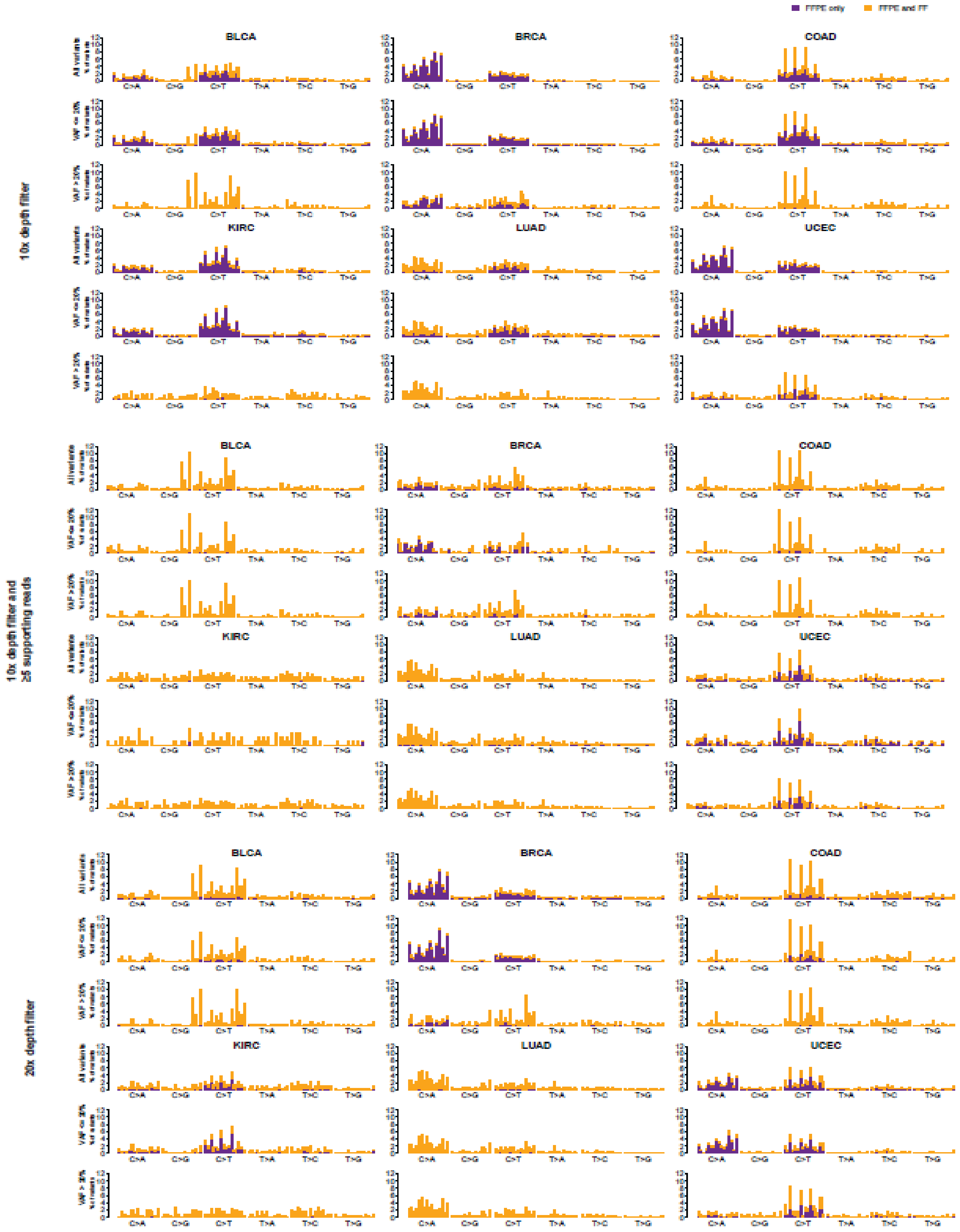

**Supplementary Figure 3**

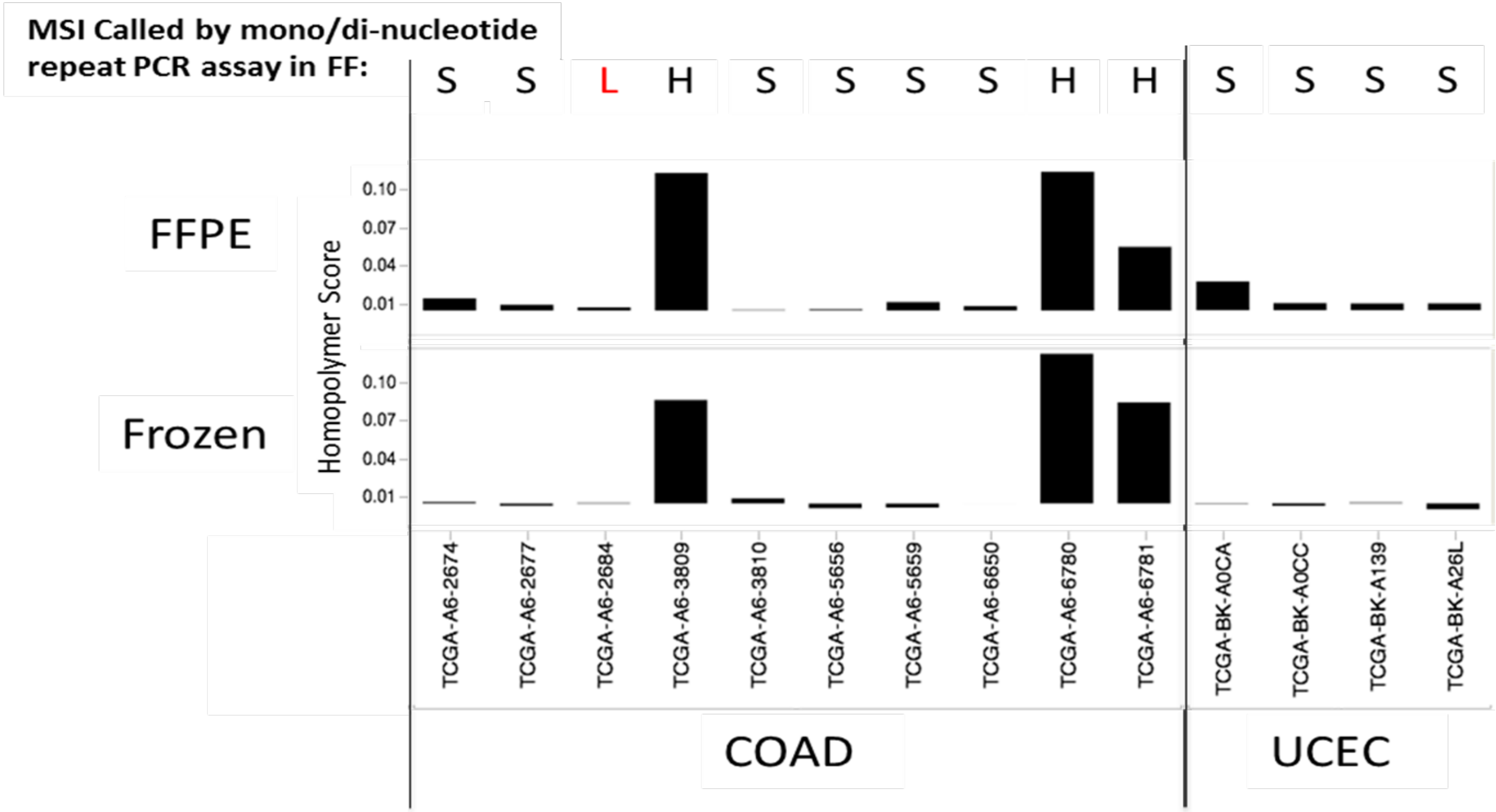

**Supplementary Figure 4**

a

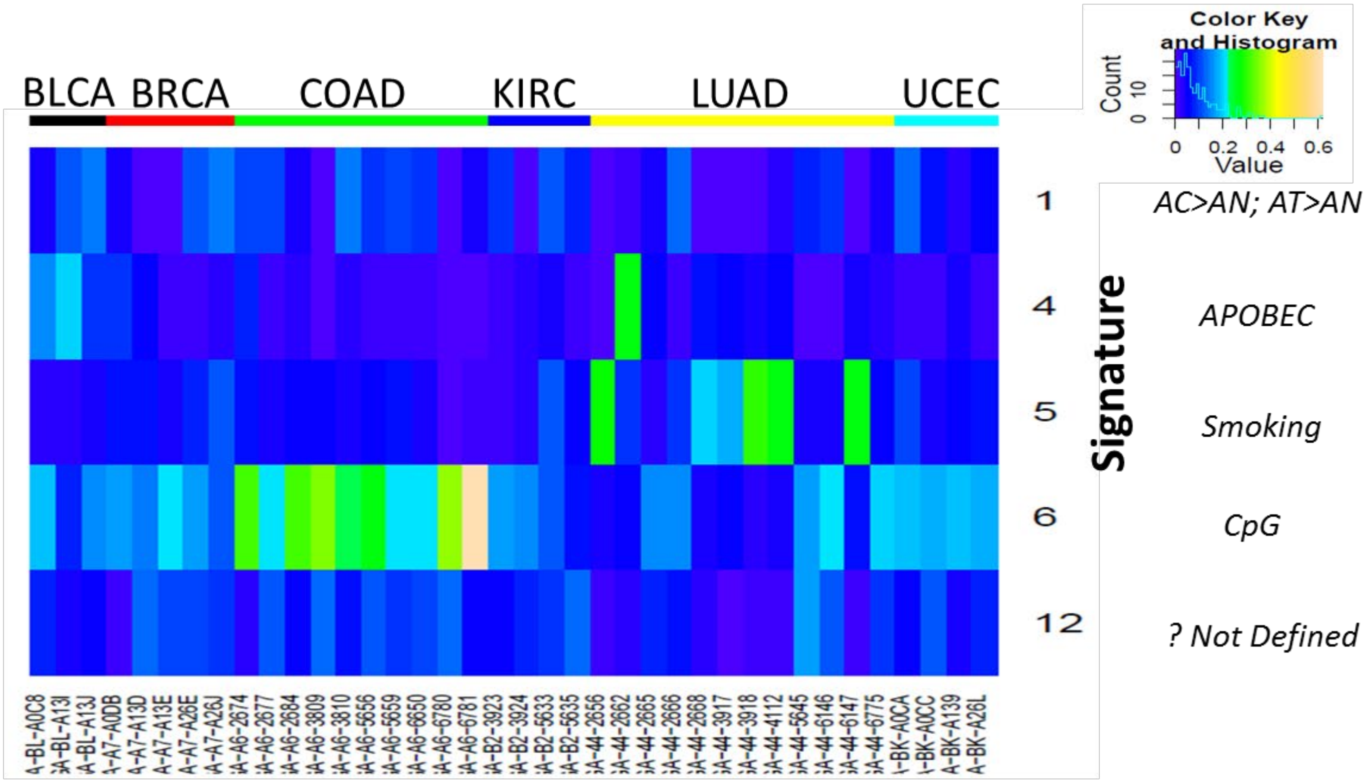


b

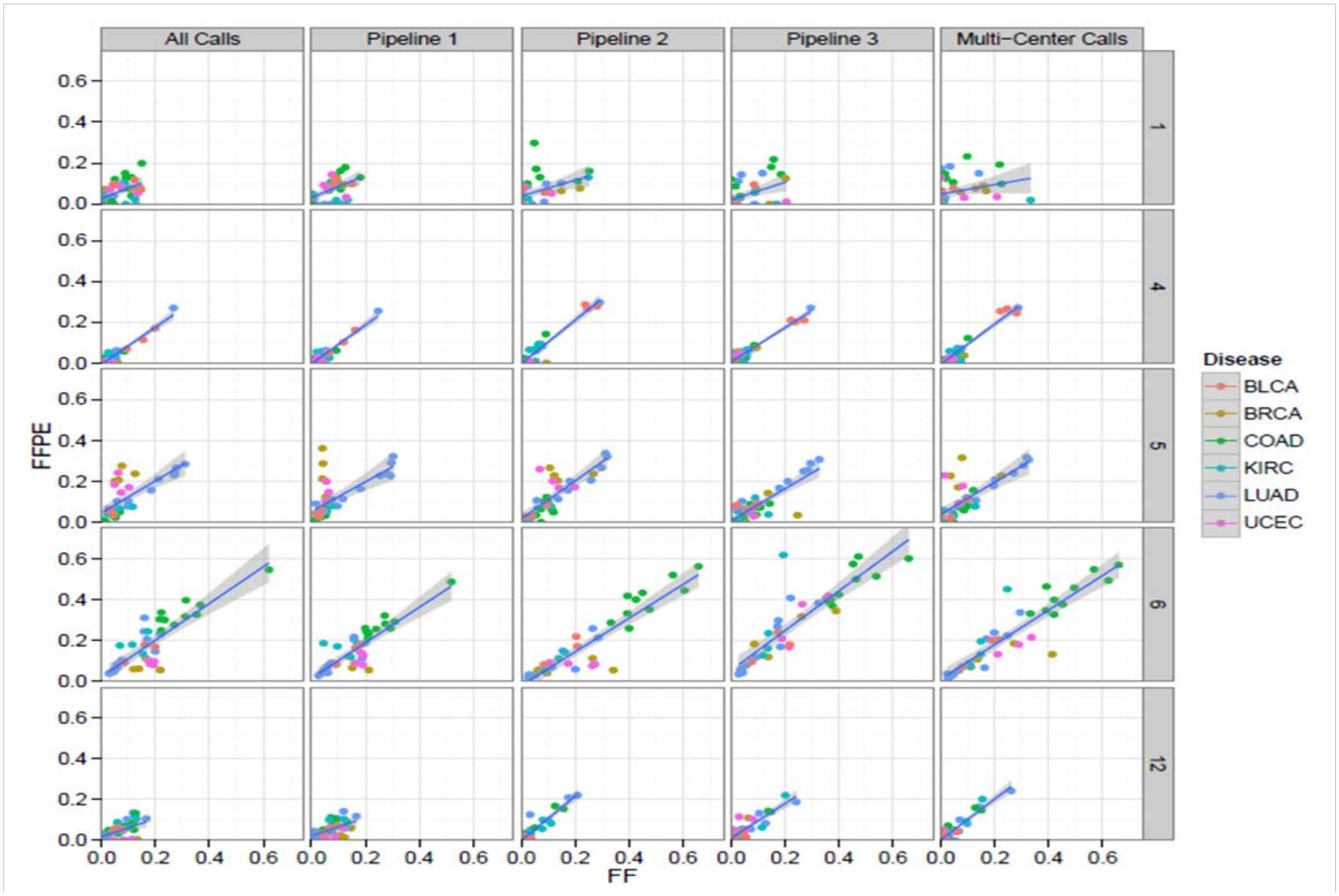

**Supplementary Figure 5**

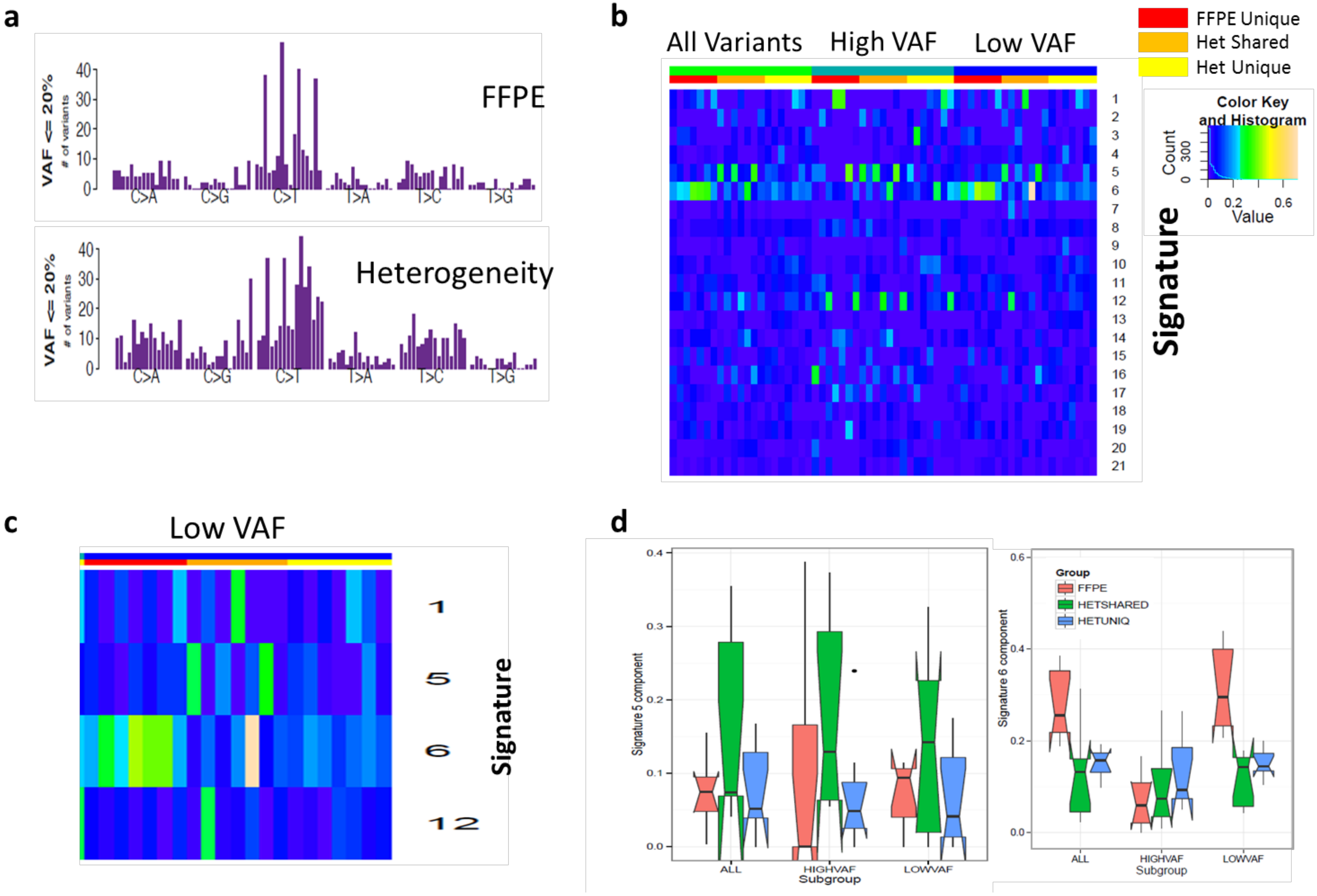

**Supplementary Figure 6**

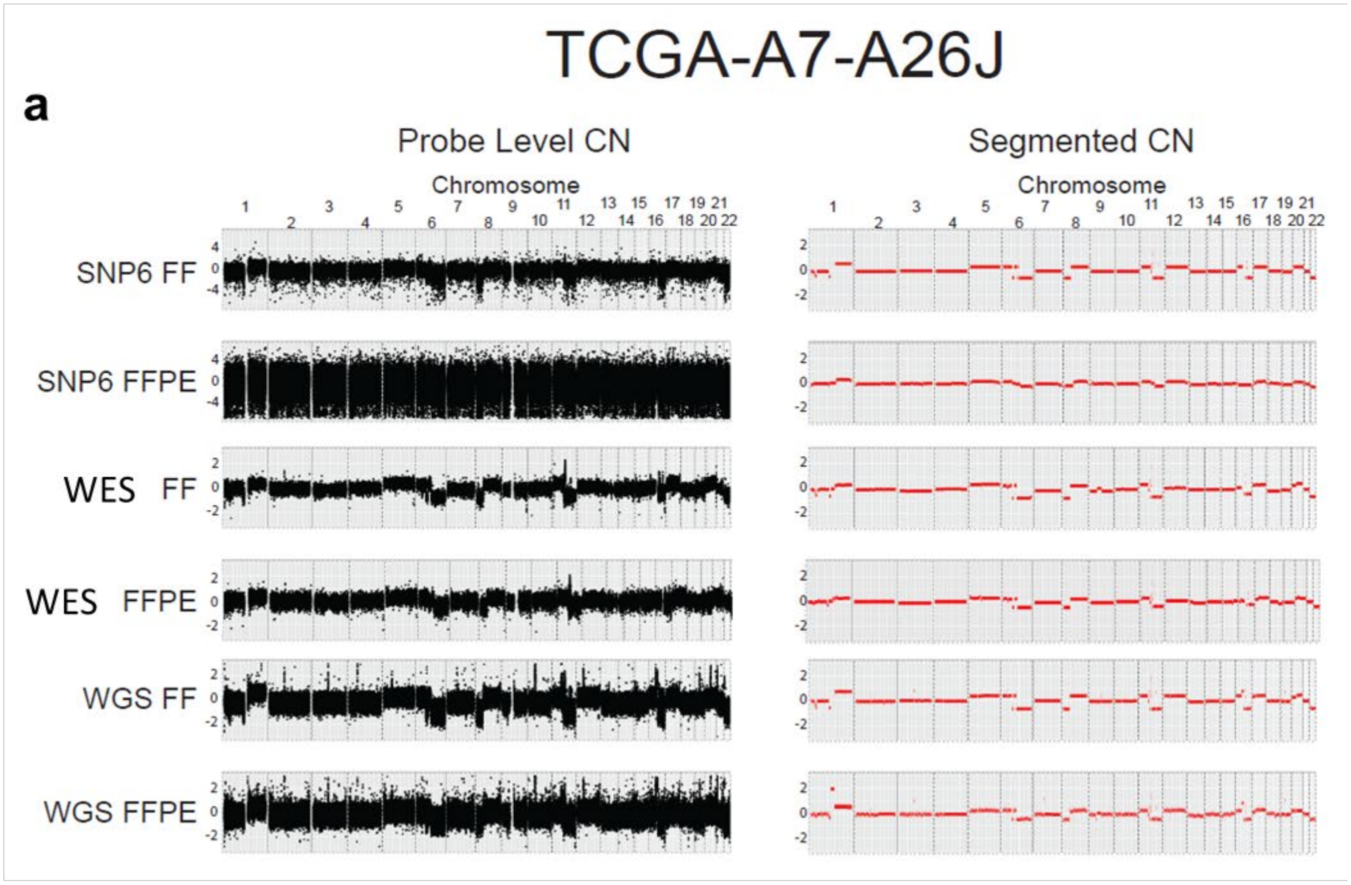


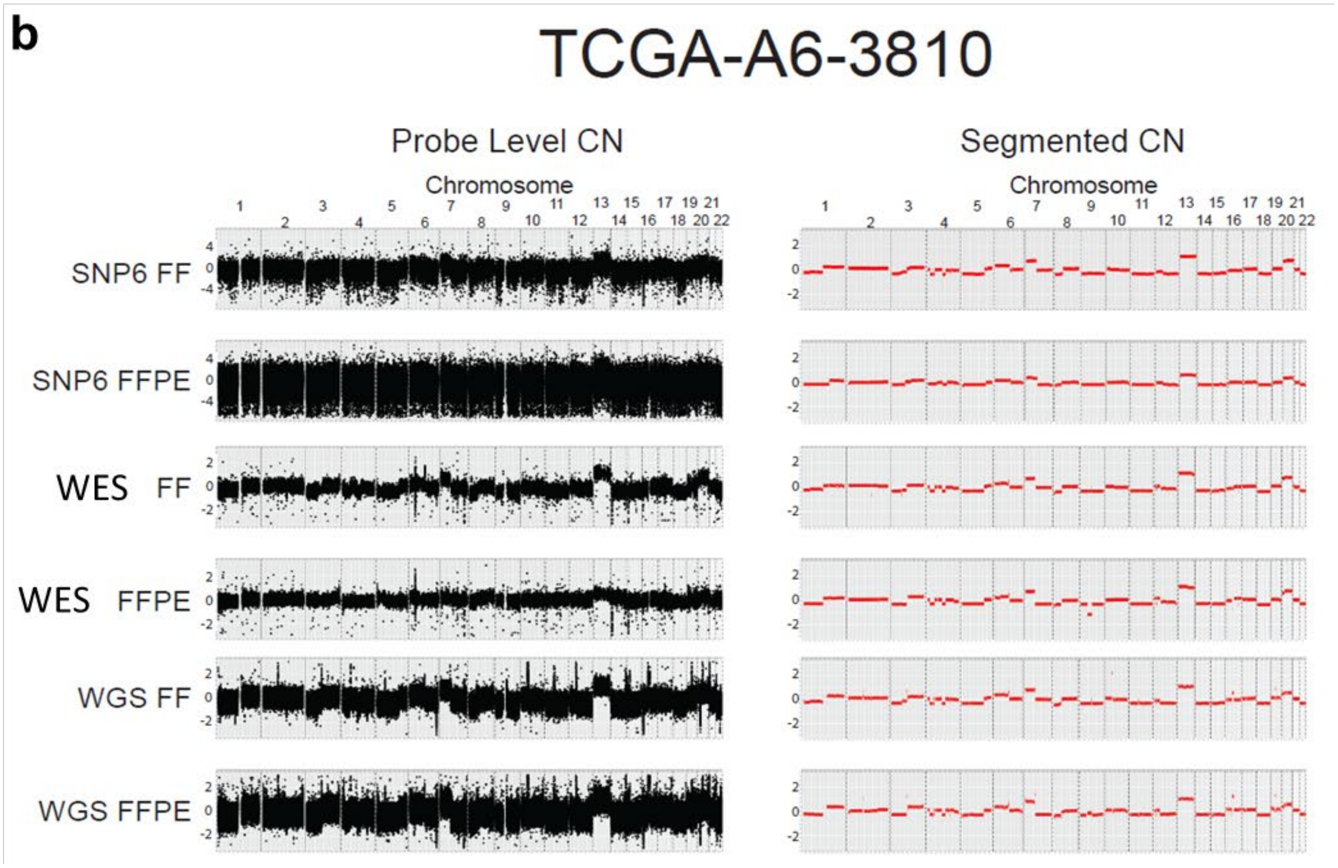

**Supplementary Figure 7**

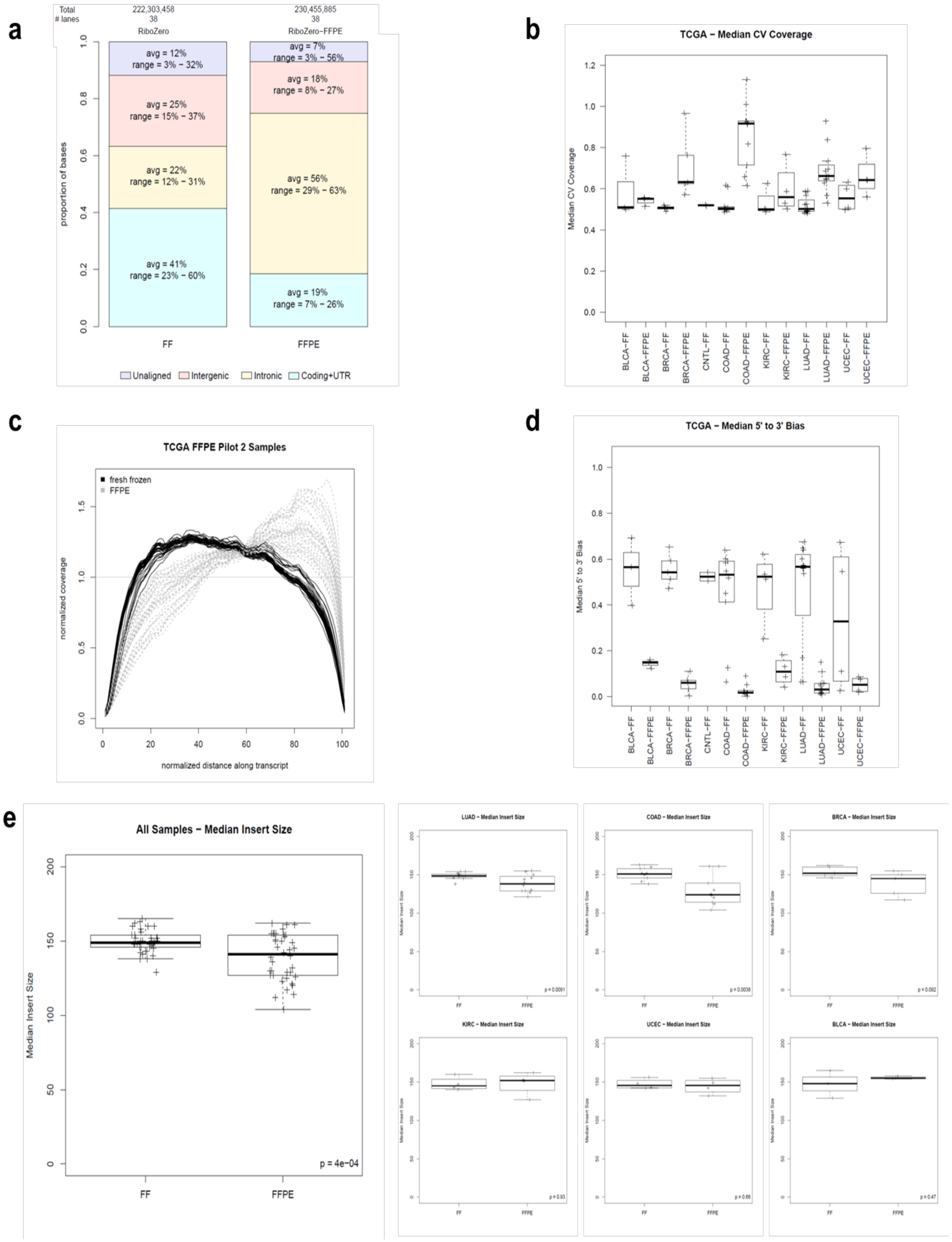

**Supplementary Figure 8**

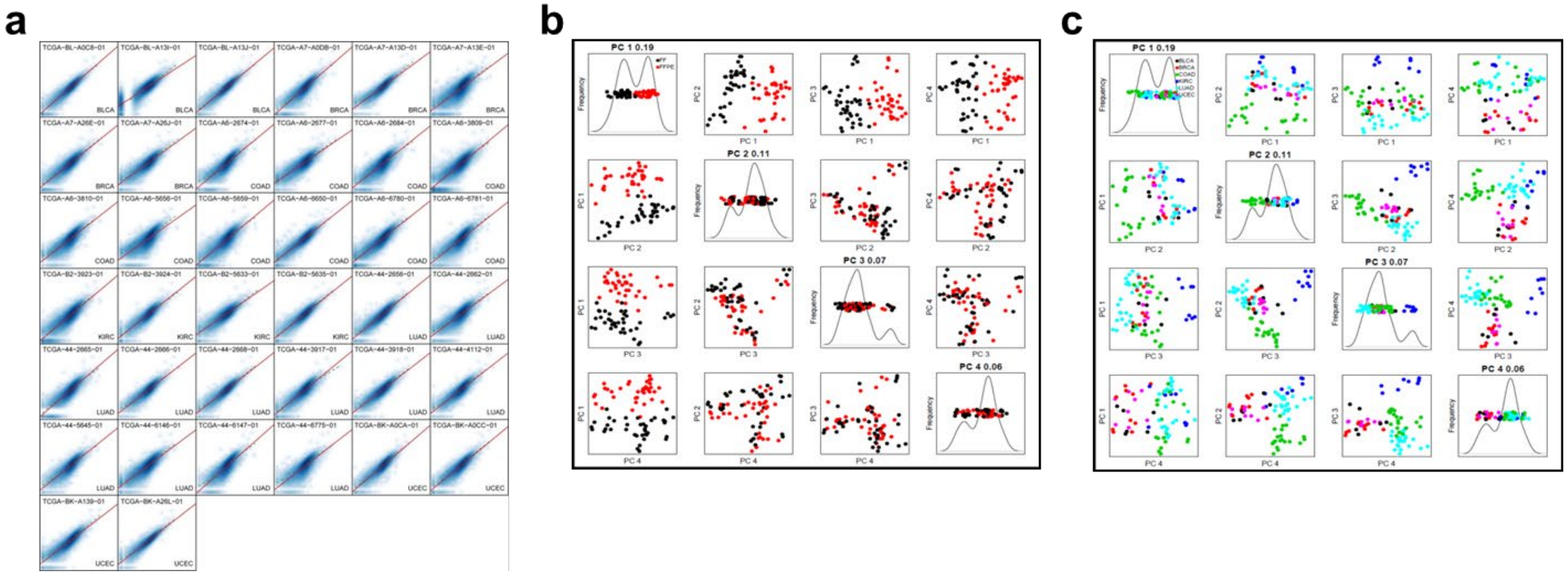

Supplementary Figure 9

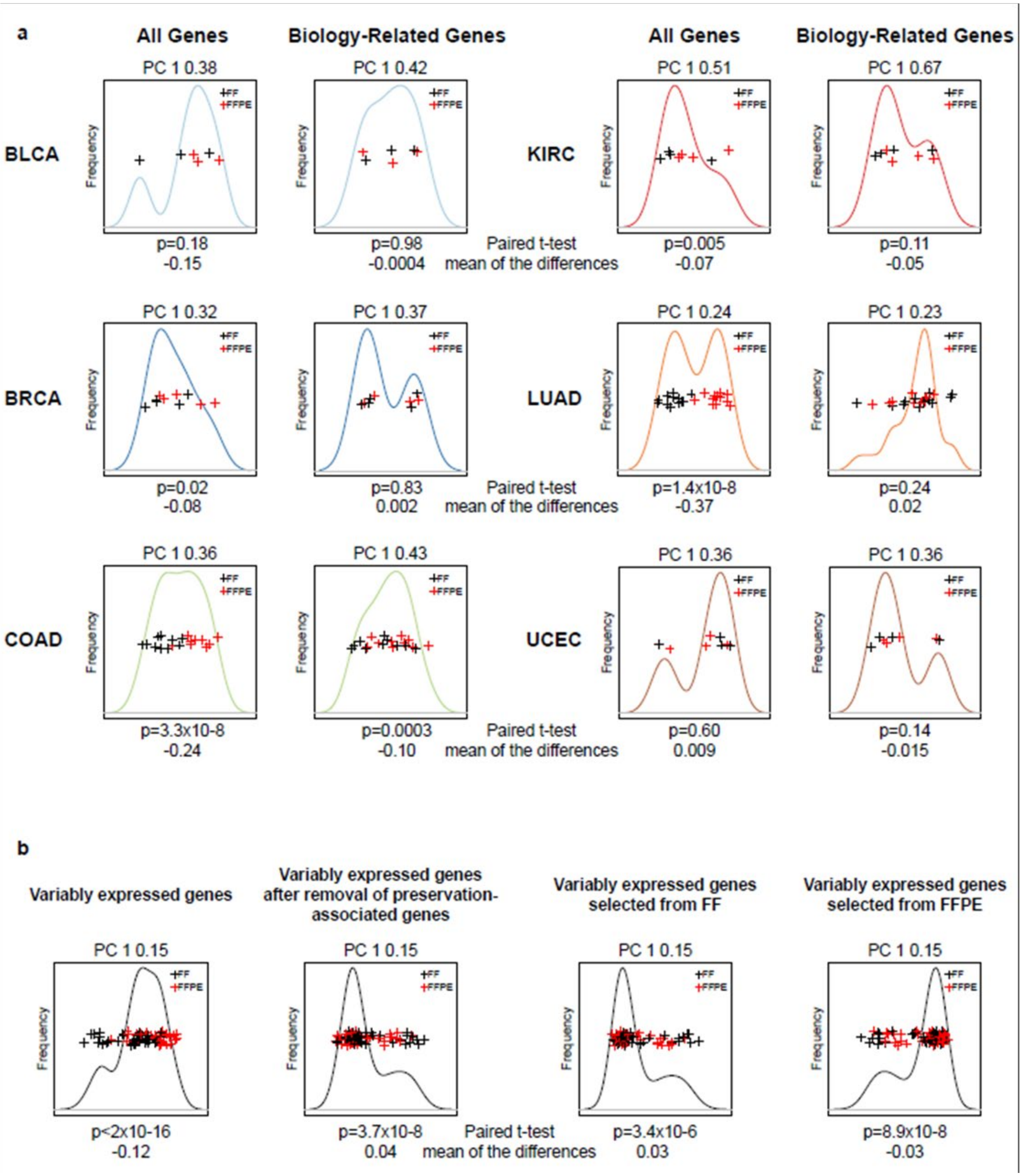

**Supplementary Figure 10**

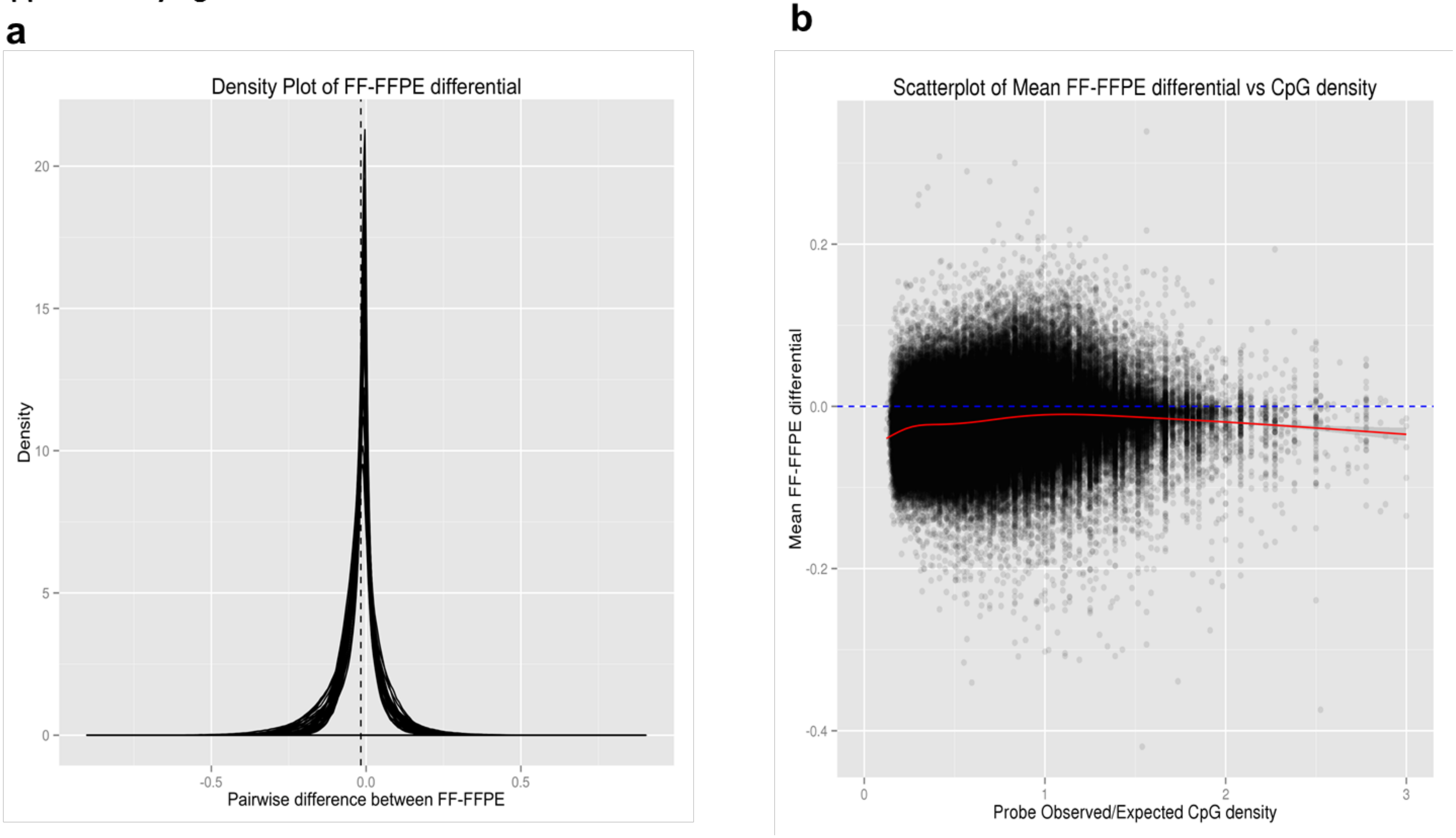

**Supplementary Figure 11**

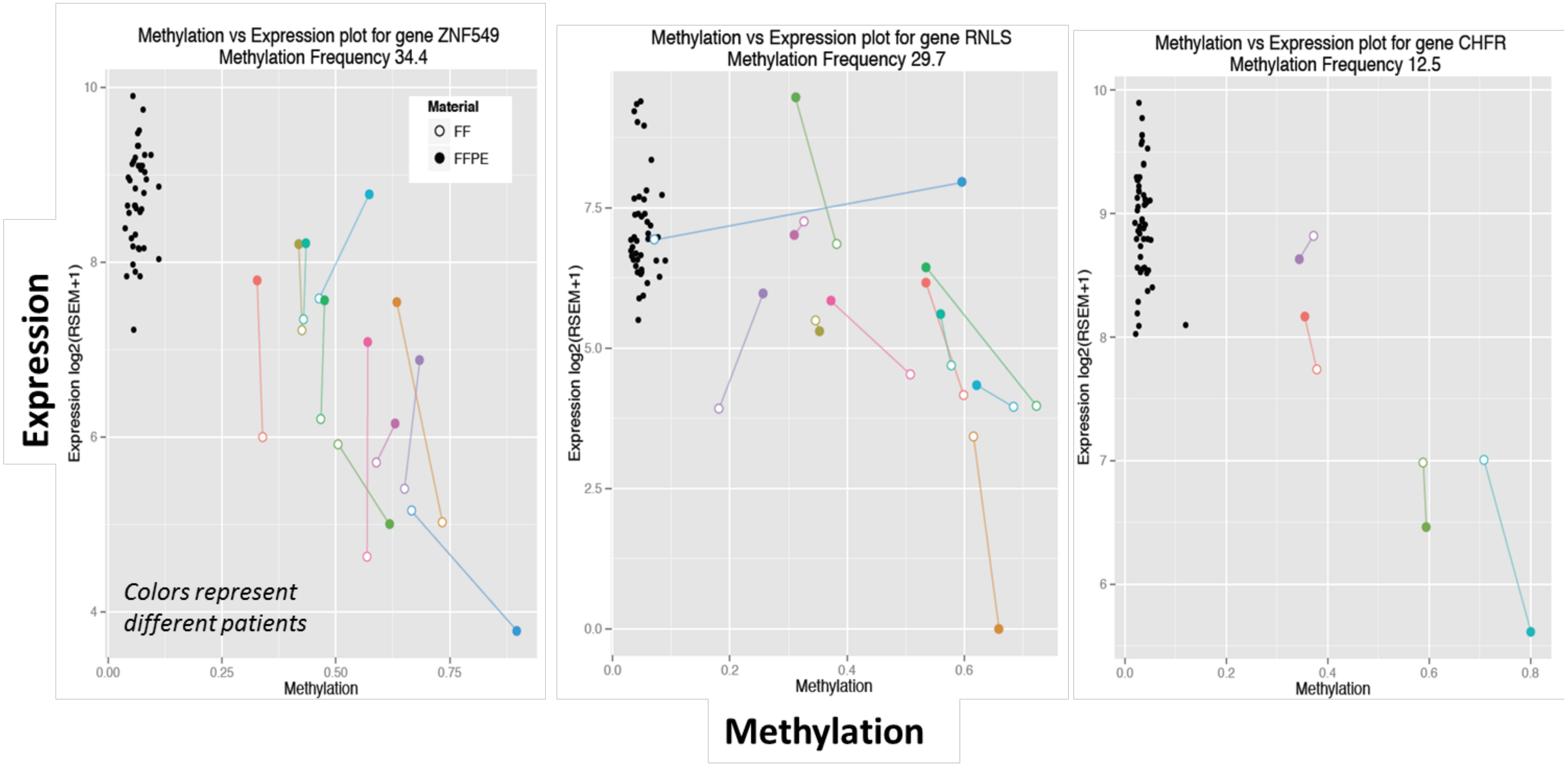

**Supplementary Figure 12**

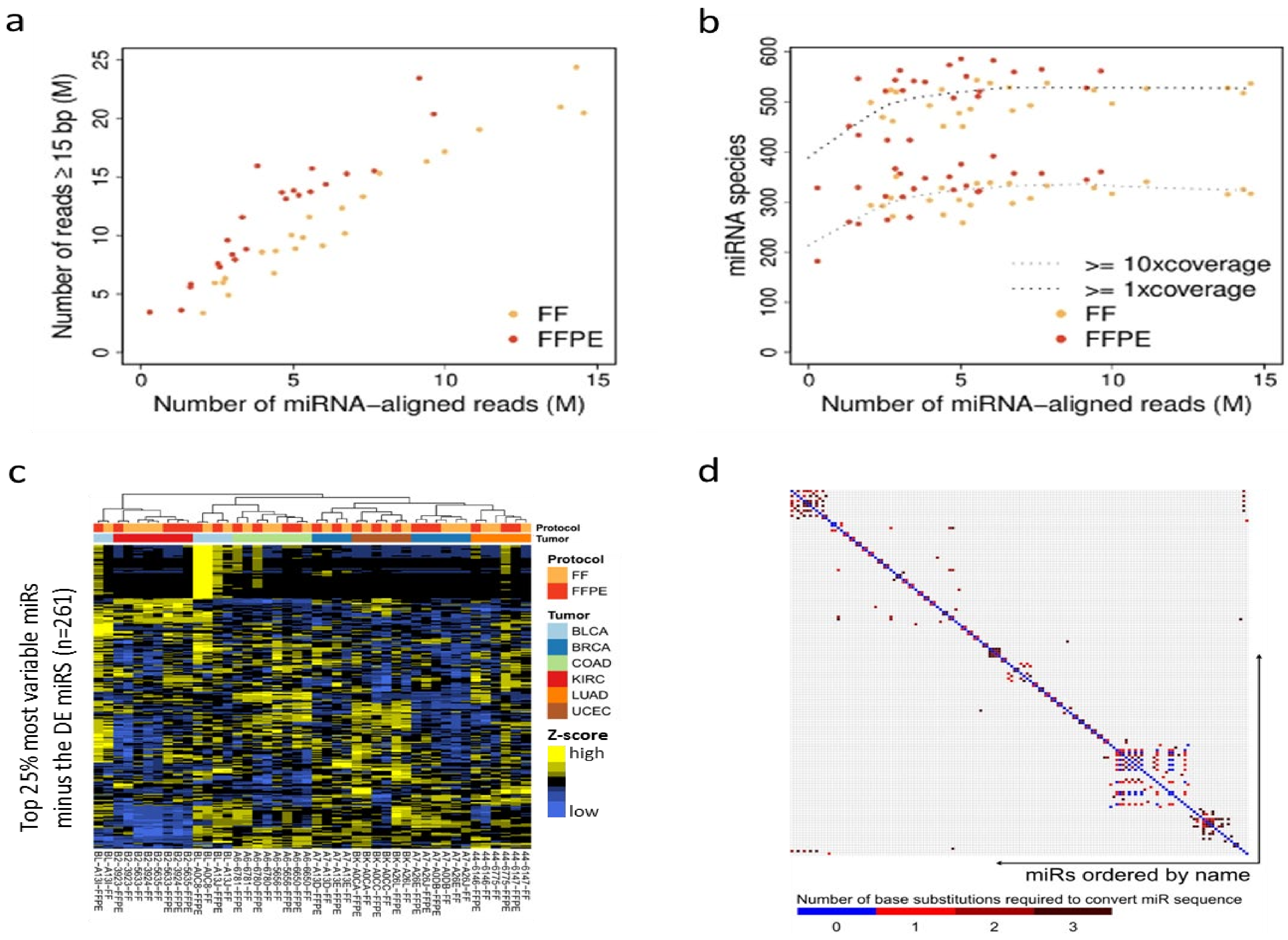

**Supplementary Fig. 13**

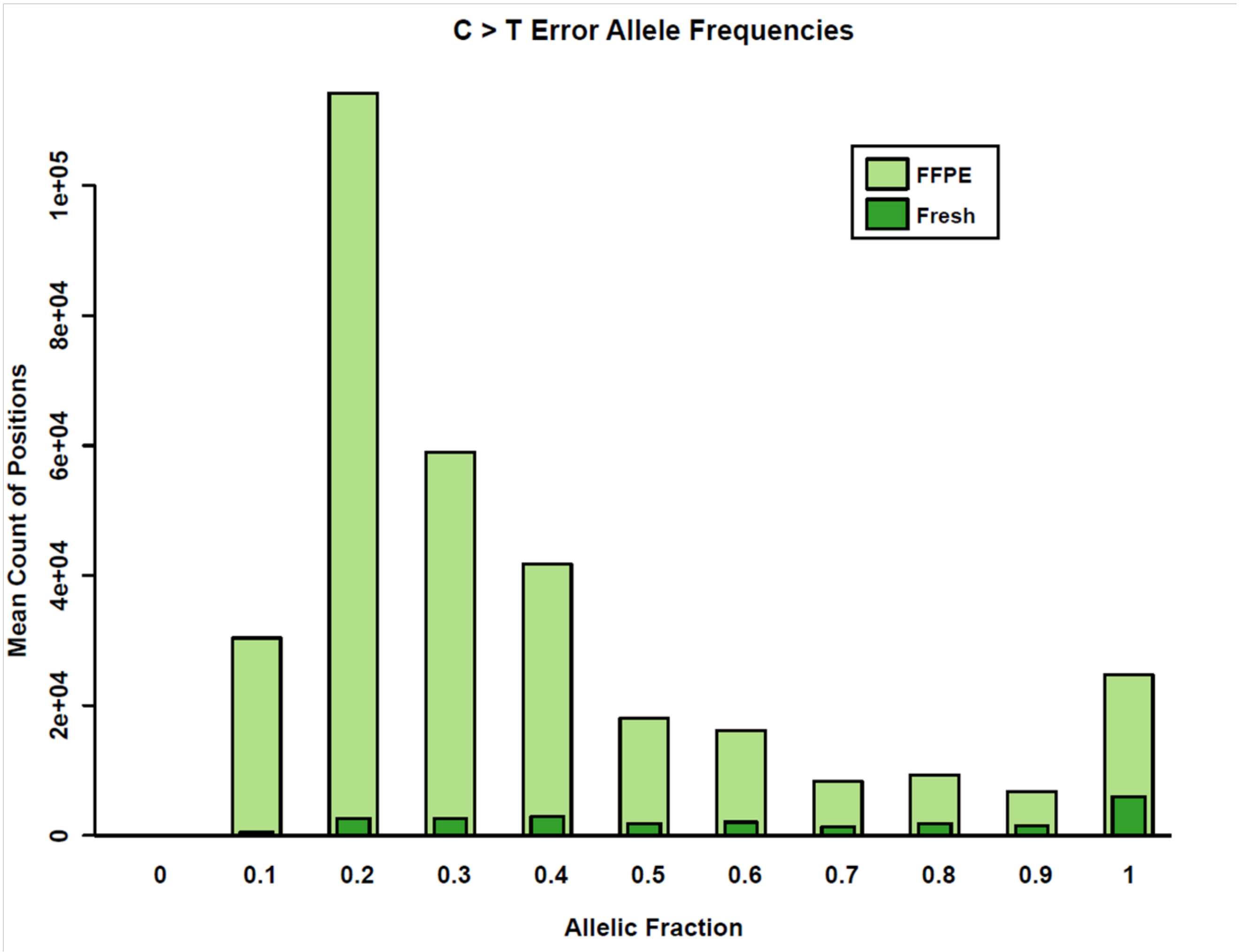
